# Inferring Progressive Disconnection in Alzheimer’s Disease with Probabilistic Boolean Networks

**DOI:** 10.1101/2025.07.10.664143

**Authors:** Liu Zhonglin, Zhang Louxin, Ching Wai Ki

## Abstract

The modern understanding of Alzheimer’s disease as a disconnection syndrome presents the challenge of quantifying the directed influence between brain regions. To address this, we apply probabilistic Boolean networks to model effective brain connectivity for the first time, introducing a novel framework for analyzing functional magnetic resonance imaging data from a cohort comprising normal controls, individuals with mild cognitive impairment (MCI), and Alzheimer’s patients. Our robust statistical analysis identified five significant connections, each exhibiting a linear decline in influence throughout the disease spectrum. We observed a progressive disruption of pathways from the Default Mode Network to the Medial Temporal Lobe, capturing a key psychophysiological mechanism underlying Alzheimer’s disease. These findings demonstrate the potential of our framework as a powerful tool for modeling network-level dynamics in neurodegeneration.

## 1 INTRODUCTION

Alzheimer’s disease (AD) is a devastating neurodegenerative disorder characterized by progressive cognitive decline. The modern understanding of AD has evolved from a focus on isolated regional pathology to a network-based perspective, viewing the disease as a “disconnection syndrome” [9]. From this viewpoint, cognitive symptoms arise from disrupted communication within and between large-scale brain networks. Functional Magnetic Resonance Imaging (fMRI) technology has been central to this shift, enabling in-vivo investigation of functional connectivity–typically defined as the statistical correlation between remote neurophysiological events [15].

Characterizing effective brain connectivity, which represents the directional neural influence between brain regions, has become increasingly important for understanding brain function and its deterioration in AD. While correlational methods provide valuable insights, they are inherently undirected and cannot reveal the causal mechanisms driving network dysfunction. To address this limitation, researchers have increasingly turned to modeling effective connectivity, defined as the directed, causal influence that one neural system exerts over another [14].

Various methodologies have been proposed for fMRI connectivity analysis, including structural equation modeling (SEM) [22], multivariate autoregressive models (MAR) [18], dynamic causal modeling (DCM) [16], and dynamic Bayesian networks (DBNs) [19], each with distinct strengths and limitations. SEM requires substantial sample sizes and assumes multivariate normality [3]. MAR models are constrained to linear interactions, inadequately capturing the nonlinear nature of neural processes. DCM encounters significant computational challenges with exponential model complexity growth [32].

DBNs have gained prominence in neuroimaging connectivity analysis [19, 31] owing to their robust statistical foundation and capacity to handle both discrete and continuous variables. DBNs can theoretically accommodate stochastic and nonlinear data characteristics while managing missing data and incorporating prior knowledge [13]. However, critical limitations make DBNs suboptimal for fMRI analysis. Practical DBN implementations assume linear Gaussian relationships due to computational constraints, limiting their ability to capture complex logical relationships. DBNs suffer from exponential computational growth with network size and require extensive temporal data for reliable parameter estimation, yet fMRI provides relatively few time points [26].

To overcome these fundamental challenges, we introduce a novel framework applying Probabilistic Boolean Networks (PBNs) [30] to fMRI-based connectivity analysis in AD. PBNs address each DBN limitation directly: they maintain computational efficiency through Boolean discretization while preserving the ability to model complex, nonlinear logical relationships between brain regions. The discrete state space enables tractable inference with limited temporal resolution typical of fMRI data, and PBNs capture switch-like, threshold-based dynamics fundamental to neural processing but lost in continuous probabilistic models. PBNs integrate logical rule-based frameworks with probabilistic characteristics, establishing them as powerful tools for systems-level analysis with computational advantages enabling practical inference of nonlinear relationships.

The clinical and translational implications of effective connectivity biomarkers are profound. Current diagnostic approaches often detect pathology only after substantial neurodegeneration has occurred. Effective connectivity measures offer potential for earlier detection of network dysfunction, crucial for early intervention strategies and disease-modifying therapies that may be most effective in preclinical stages. Quantitative connectivity biomarkers could transform clinical trial design by providing objective end-points that capture disease progression with greater sensitivity than traditional cognitive measures, enabling smaller trials while providing mechanistic insights into therapeutic effects. Clinical applications have validated PBN effectiveness in Parkinson’s disease, demonstrating detection of pathological network changes and therapeutic monitoring capabilities [21].

However, applying PBN to AD fMRI analysis requires novel approaches for signal binarization in resting-state data and integration of prior knowledge given limited temporal data. We develop a novel PBN inference framework for resting-state fMRI data spanning the complete AD spectrum: cognitively normal controls, individuals with MCI, and patients with diagnosed AD. Our methodology incorporates a hybrid structure learning algorithm integrating data-driven dynamics with neuroanatomical priors and specialized pre-processing techniques. We focus on vulnerable networks including the Default Mode Network (DMN), Executive Control Network (ECN), Salience Network (SN), and Medial Temporal Lobe (MTL) network, particularly examining DMN-MTL connectivity given established pathological vulnerabilities.

We hypothesize that PBN approaches can reliably detect and quantify progressive neural pathway degradation across the AD continuum with superior computational efficiency and interpretability compared to DBN methods. Key findings demonstrate systematic linear decline in directed DMN-to-MTL influence across disease stages, providing quantitative evidence for fundamental AD pathophysiological mechanisms with immediate translational relevance for biomarker development and therapeutic monitoring.

The remainder of this paper is organized as follows. Section 2 describes our PBN framework for effective connectivity analysis. Section 3 presents comprehensive analysis across the AD spectrum. Section 4 discusses results, neurobiological interpretation, and clinical implications.

## 2 METHODS

### 2.1 Participant Data and fMRI Preprocessing

All data used in this study were obtained from the Alzheimer’s Disease Neuroimaging Initiative (ADNI) database^1^. For our analysis, we specifically utilized the preprocessed resting-state fMRI series labeled as “Axial MB rsfMRI (Eyes Open)”.

Participants were assigned to one of three groups based on a comprehensive set of diagnostic and biomarker criteria:

- Normal Control (NC, n=21),
- Mild Cognitive Impairment (MCI, n=28), and
- AD (n=10).

The NC group consisted of cognitively normal individuals with a Mini-Mental State Examination (MMSE) score between 28-30, a Clinical Dementia Rating (CDR) Global Score of 0.0 or 0.5, and a CDR Memory score of 0.0. A key inclusion criterion for this group was a “Non-Elevated” status on amyloid Positron Emission Tomography (PET) scans, indicating no evidence of significant cortical amyloid-beta plaque deposition.

The AD group included individuals with a clinical diagnosis of Dementia, characterized by MMSE scores in the range of 0-23 and significant functional and memory impairment (CDR Global Scores of 0.5-3.0; CDR Memory scores of 1.0-3.0). Critically, these participants were required to have an “Elevated” amyloid-PET status, which provides direct *in vivo* evidence of amyloid pathology, a core neuropathological hallmark of AD and a key component of modern biological definitions of the disease [20].

The MCI group, represents a transitional stage. The group included participants with MMSE scores between 24-27 and a CDR Global Score of 0.5. To specifically select for an amnestic MCI subtype typical of prodromal AD, a key criterion was a score between 0.0-7.0 on the Logical Memory Delayed Recall test. The test is a sensitive measure of episodic memory, and impairment on this test is a well-established indicator of the memory deficits characteristic of early-stage AD [35].

Functional and anatomical data were preprocessed using fMRIPrep 25.1.1 [10], a comprehensive neuroimaging pipeline based on Nipype 1.8.6 [17]. The T1-weighted anatomical image for each subject underwent intensity non-uniformity correction with `N4BiasFieldCorrection` [33], brain extraction, and cortical surface reconstruction using `recon-all` from FreeSurfer v7.3.2 [8]. The T1-weighted reference was then spatially normalized to the `MNI152NLin2009cAsym` template [11] through nonlinear registration with `antsRegistration` [1]. The Blood-Oxygen-Level-Dependent (BOLD) data were corrected for susceptibility distortion, slice-timing using `3dTshift` from AFNI [7], and head motion using `mcflirt` (FSL 6.0.7.1). The corrected BOLD series was then co-registered to the subject’s T1-weighted reference and spatially normalized to the standard template in a single interpolation step.

### 2.2 ROI Definition and Binarization

To investigate functional connectivity alterations in AD, we defined 18 regions of interest (ROIs) based on the Automated Anatomical Labeling 3 atlas [27]. These ROIs were selected as key nodes within four large-scale brain networks known to be highly vulnerable to AD pathology. The DMN, critical for episodic memory, is a primary site of early amyloid-*β* deposition. The Executive Control Network (ECN) is crucial for higher-order cognitive functions that are progressively impaired in AD. The SN mediates cognitive control, and its dysfunction contributes to attentional deficits. Finally, the MTL network is foundational for memory and among the first to exhibit severe atrophy. Our selection comprised the bilateral Precuneus (PREC), Angular Gyrus (ANG), and Medial Orbital Frontal Gyrus (FMO) for the DMN (6 ROIs); the bilateral Superior Frontal Gyrus (FSG) and Superior Parietal Gyrus (PARS) for the ECN (4 ROIs); the bilateral Insula (INS) and Supplementary Motor Area (SMA) for the SN (4 ROIs); and the bilateral Hippocampus (HIPP) and Parahippocampal Gyrus (PHC) for the MTL (4 ROIs). A complete mapping of all 18 ROIs with their abbreviations and network assignments can be found in Appendix B.

The extraction and processing of the BOLD signal for the ROIs followed a multi-step procedure. All preprocessed fMRI data were first harmonized to a repetition time (TR) of 0.61 seconds. Using Statistical Parametric Mapping 12 (SPM12), a binary mask for each of the 18 ROIs was generated from the Automated Anatomical Labeling 3 (AAL3) atlas, resampled into each participant’s native functional space, and used to extract the mean BOLD time series. To prepare the data for PBN analysis, the continuous signals were denoised using linear detrending and a bandpass filter (0.01–0.1 Hz) to remove scanner drift and isolate neurally relevant frequencies.

The crucial final step was the binarization of the denoised time series using a Hidden Markov Model (HMM) [25]. Our approach was inspired by the work of Ma et al. [21] but adapted for resting-state fMRI. Their method leveraged known task timings to inform state transitions; the absence of such a structure in our data necessitated the development of a novel, fully data-driven iterative HMM. This model was designed to infer the transitions between unobserved ‘low’ and ‘high’ neural activity states directly from the temporal dynamics of the signal itself.

Mathematically, the HMM is defined by *λ* = (*S, O, A, B, π*). For our application, *S* represents the two hidden states, and *O* is the observed BOLD signal. The dynamics are governed by the state transition matrix *A*, initialized with a higher probability of remaining in the same state (*a*_*ii*_ = 0.85), and the observation probability distribution *B*, which we model as a Gaussian:

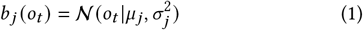

where *μ* _*j*_ and 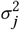 are the mean and variance of state *s* _*j*_. Our implementation begins by initializing these state parameters using K-Means clustering. The model is then iteratively refined over three cycles using the Viterbi algorithm [12] to infer the most probable sequence of hidden states, followed by a re-estimation of the state parameters. A final post-processing step ensures temporal stability by merging any state lasting less than 3 TRs. This procedure yields a plausible binary time series for each ROI, as shown in Fig. 1, ready for PBN inference.

**Figure 1:**
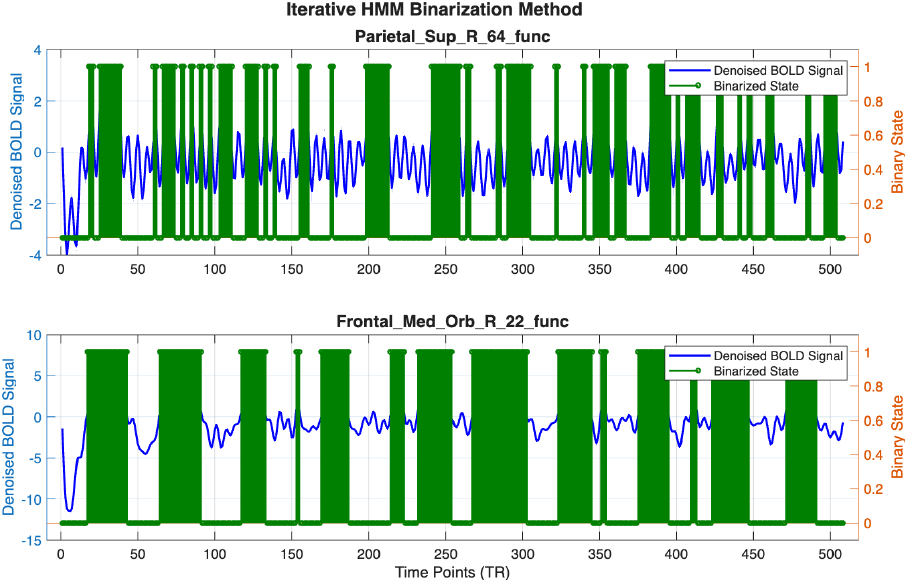
Example output of the iterative HMM binarization process, demonstrating its behavior on two distinct signals from a representative subject. The signal for ROI 1 (gray) exhibits low variance and is correctly classified as the ‘low’ state throughout. In contrast, the signal for ROI 2 (blue) shows significant fluctuation. The corresponding binary state sequence (red stems) successfully captures these dynamics, switching between ‘low’ (0) and ‘high’ (1) states. This illustrates the robustness of the algorithm in handling different levels of neural activity.

### 2.3 PBNs

#### 2.3.1 Definition and Properties

PBNs represent a significant advancement over traditional BNs by incorporating stochastic properties to model complex interactions among different nodes within a system. A BN consists of a set of nodes *V* = {*v*_1_, *v*_2_, …, *v*_*n*_} and a corresponding list of Boolean functions *F* = {*f*_1_, *f*_2_, …, *f*_*n*_}, where each Boolean function *f*_*i*_ is defined as a combination of logical gates (AND, OR, NOT) that regulate the connections among nodes. In this deterministic framework, each node exhibits switch-like behavior, transitioning between on and off states in the temporal dimension, with the activity of each node being governed by parent nodes through a single, fixed Boolean function. Values of nodes are synchronously updated by corresponding functions based on the current state of parent nodes, which serve as input variables to these functions.

The fundamental limitation of BNs lies in their deterministic rigidity. The temporal evolution of the entire system is completely determined by the initial state, with the system following a static transition mechanism regulated by binary functions and ultimately converging to an attractor state from which it cannot move. All possible states that converge to the same attractor state are grouped into the basin of that attractor, forming distinct basins of attraction.

To address these limitations, Shmulevich et al. proposed PBNs, which accommodate multiple Boolean functions for each node, thereby providing substantially greater modeling flexibility [30]. The key distinction between PBNs and BNs lies in the probabilistic selection mechanism: while a BN employs a single, fixed Boolean function for each node, a PBN maintains a set of Boolean functions known as predictors for each node. The fundamental recurrence relation that governs PBN dynamics is expressed as:

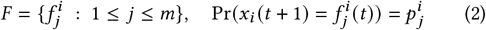

where *m* is the number of predictors for node *x*_*i*_, 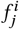 represents the *j*-th predictor function for node *i*, and 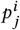 denotes the probability of selecting predictor *j* at each transition, with the constraint 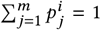. This fundamental equation captures the essence of PBN stochasticity, where at each time step, one predictor is probabilistically selected to update the target node’s state according to its associated probability.

To illustrate the conceptual difference, consider extending a three-node BN example to a PBN framework. In the original BN, node *v*_1_ might be governed by a single Boolean function *f*_1_ (*v*_2_, *v*_3_) = *v*_2_ ∧ ¬*v*_3_. In the corresponding PBN, node *v*_1_ could have multiple predictors: 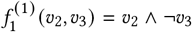 with probability 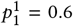 and 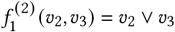 with probability 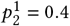. At each time step, the system probabilistically selects between these two functions according to Equation (2) to determine the next state of *v*_1_, introducing stochastic variability that captures the inherent uncertainty in biological systems.

The dynamic behavior of a PBN is characterized by a Markov chain whose transition matrix is defined by all possible Boolean functions and their corresponding probabilities. This probabilistic framework enables the system to exhibit more realistic behaviors compared to the rigid deterministic dynamics of traditional BNs. The long-run asymptotic behavior of the PBN is of particular interest, specifically whether there exists a steady-state distribution that the system approaches regardless of its initial distribution. The system’s reducibility properties determine whether it possesses single or multiple stationary distributions. An irreducible PBN, characterized by a communicating state space where any state can potentially be reached from any other state, possesses a unique stationary distribution. In contrast, a reducible PBN contains irreducible sets of states, with each set maintaining its own unique stationary distribution.

The steady-state behavior can be formally analyzed through the stationary distribution *π*, defined such that the probability of the system being in state *i* at any time *t* equals *π*_*i*_ when the initial distribution is *π*. A PBN achieves a steady-state distribution if and only if it is both irreducible and aperiodic (ergodic), satisfying the condition:

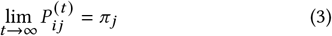

where 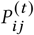 represents the *t*-time transition probability from state *i* to state *j*. This probabilistic framework allows PBNs to capture the stochastic nature of biological networks while maintaining the interpretability and computational efficiency of Boolean logic.

#### 2.3.2 Learning PBNs

The inference of network models from fMRI time series is an inherently ill-posed problem, as the number of potential interactions far exceeds the number of available time points. Methods that rely solely on time-series data are therefore susceptible to overfitting and may produce biologically implausible network structures. To overcome this challenge, we developed a multi-stage inference pipeline inspired by recent structure-aware methodologies like SAILOR [24]. Our framework is designed to systematically determine the optimal set of predictive Boolean functions for each node by synergistically integrating data-driven evidence from fMRI time series with neurobiological priors from established reference networks.

Our inference pipeline begins by constructing a single hybrid influence matrix, *W*_hybrid_, for each subject, which balances data-driven dynamics with a structural reference. To create the structural reference, we first compute a group-average functional connectivity matrix from the Pearson correlations of all subjects’ fMRI time series. From this average matrix, we generate N=10 binary reference networks by thresholding the top 5% to 50% of connection strengths. These ten networks are then averaged to form a probabilistic consensus matrix, *A*_con_, where each entry represents the connection probability based on its prevalence across the different thresholds.

Concurrently, for each subject, the continuous BOLD time series is analyzed using the `dynGENIE3` algorithm to produce an 18 × 18 data-driven influence matrix, *W*_dyn_. These two sources of information—the consensus reference (*A*_con_) and the individual’s dynamics (*W*_dyn_)—are subsequently combined into the final hybrid matrix using a weighted linear combination:

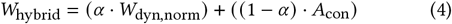

where *W*_dyn,norm_ is the *W*_dyn_ matrix scaled to a [0, 1] range, and *α* is set to 0.7. Next, to manage the immense search space of potential predictors, a two-step pruning strategy is employed. For each of the 18 ROIs, the *W*_hybrid_ matrix is used to identify the top potential regulators(set to 10 for computational efficiency) with the highest influence on that target node. From this focused subset of likely parent nodes, we then generate all possible combinations of regulators, with the size of each combination ranging from 1 to a constraint of 6. This combinatorial process yields a comprehensive yet manageable set of candidate predictor functions to be evaluated.

Following the search, for each candidate combination of parent nodes, the most parsimonious Boolean rule describing the relationship is inferred using the Quine-McCluskey algorithm on the subject’s binarized time series data. The “goodness-of-fit” for each inferred rule is then evaluated by calculating its Coefficient of Determination (COD). The COD measures the proportional reduction in prediction error achieved by the Boolean function compared to a simple constant estimator (the mean of the target node’s time series). It is defined as:

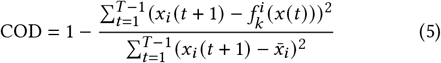

where *x*_*i*_ (*t* + 1) is the observed state of the target node *i* at time 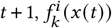 is the state predicted by the Boolean function, and 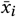 is the mean state of the target node. A higher COD score indicates superior predictive power.

Finally, after all candidate functions have been inferred and scored, they are ranked by their COD values. For each of the 18 ROIs, the top 4 predictors with the highest COD scores are selected as the final set of predictors for that node. To assign a probability to each of these selected functions, a normalization step converts the fitness scores into a true probability distribution 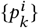 for each node *i*. The probability for the *k*-th predictor of node *i* is given by:

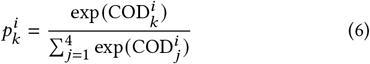

where 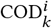 is the Coefficient of Determination for the *k*-th selected predictor. The final output of this pipeline is a comprehensive, subject-specific PBN.

Upon obtaining the set of predictors and their associated probabilities, the quantification of inter-node relationships is accomplished through influence value calculations. The influence of node *v* _*j*_ on node *v*_*i*_, denoted as *I* (*v* _*j*_ *→ v*_*i*_), is defined as the sum of influences across all predictors of *v*_*i*_, weighted by their respective probabilities:

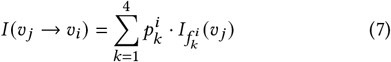

where 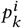 is the probability of the *k*-th predictor function for node *v*_*i*_, and 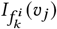 is the influence of *v* _*j*_ within that specific function.

This per-predictor influence is calculated as the sensitivity of the function’s output to the toggling of input *v* _*j*_, averaged over all possible input states:

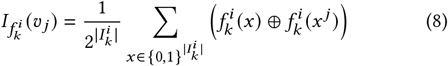

where 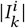 is the number of inputs to function 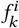, *x* ^*j*^ represents the input vector *x* with the value of *v* flipped, and ⊕ denotes the exclusive OR operation. This calculation is applied across all 18 ROIs, resulting in an 18 × 18 influence matrix that serves as a weighted directed graph of the subject’s brain network. An example of such a network is visualized in Fig. 2.

**Figure 2:**
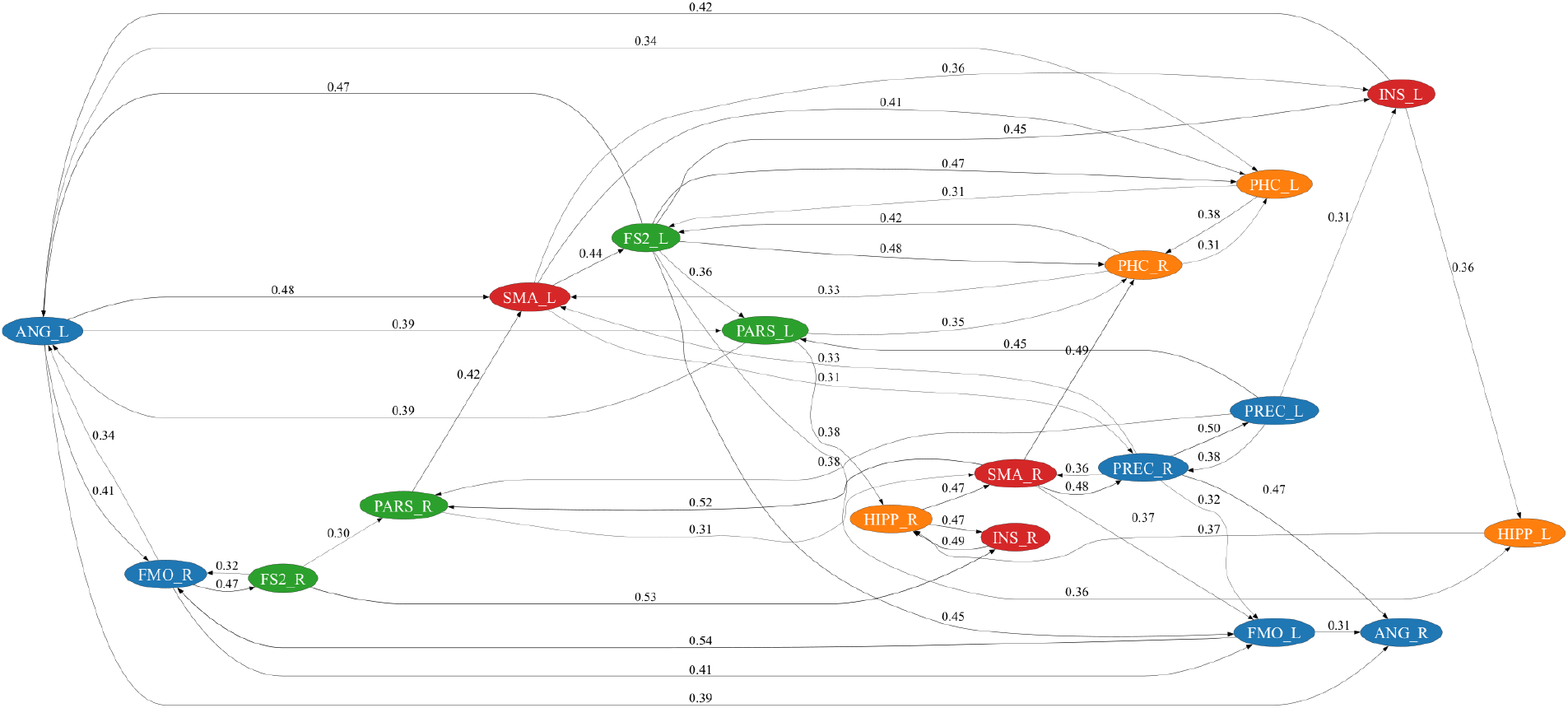
An example of a learned influence network for a single subject. Nodes are colored by their respective large-scale brain networks: Default Mode Network (DMN, blue), Medial Temporal Lobe (MTL, orange), Executive Control Network (ECN, green), and Salience Network (SN, red). Edges represent the directed influence from one ROI to another, with arrow thickness corresponding to the influence magnitude. Only connections above an influence threshold of 0.3 are shown for clarity.

## 3 RESULTS

The PBN inference framework detailed in the preceding section was applied to the fMRI data of each participant in the NC, MCI, and AD cohorts. This process yielded a subject-specific 18 × 18 influence matrix for each individual, quantifying the directed connectivity between the selected brain regions. These matrices were then carried forward for group-level statistical analysis to identify systematic alterations in network connectivity across the AD continuum. This section presents the results of this analysis and discusses their neurobiological implications.

### 3.1 Statistical Analysis

To address inter-subject variability and identify significant group-level differences, a mass-univariate statistical approach was employed. For each of the 18 × 17 = 306 possible directed connections in the influence matrices, we tested for a significant effect of diagnostic group (NC, MCI, AD). Given the unbalanced group sizes, a Levene’s test for homogeneity of variances was first performed. For connections where the assumption of equal variances was met (*p >*.05), a standard one-way Analysis of Variance (ANOVA) was used. If the assumption was violated, a Welch’s ANOVA, which is robust to heteroscedasticity, was performed [36].

To correct for the large number of simultaneous comparisons, the resulting p-values were adjusted using the Benjamini-Hochberg False Discovery Rate (FDR) procedure [2]. A connection was considered statistically significant if its FDR-corrected q-value was less than 0.05. For these significant connections, post-hoc pairwise comparisons were conducted to determine which specific group pairs differed, using either a Tukey’s HSD test or a Games-Howell test, corresponding to the initial choice of ANOVA or Welch’s ANOVA, respectively.

### 3.2 Significant Connectivity Changes in AD

The analysis revealed 5 connections that exhibited a statistically significant difference between the groups after FDR correction. A subsequent disease progression analysis was performed to characterize the pattern of change across the clinical stages (Normal → MCI → AD). As illustrated in Fig. 5, all 5 significant connections displayed a linear decrease pattern, where the influence value was highest in the Normal group, intermediate in the MCI group, and lowest in the AD group.

The primary finding is a profound, progressive disconnection of pathways originating from the DMN and Executive Control Network (ECN) that target the MTL memory system. The DMN, critical for episodic memory, includes regions like the Precuneus, Angular Gyrus, and Medial Orbital Frontal Gyrus. The MTL is foundational for memory and includes the Hippocampus and Parahippocampal Gyrus. The results are summarized in Table 1 and visualized in Fig. 3, Fig. 4 and Fig. 6.

**Table 1:**
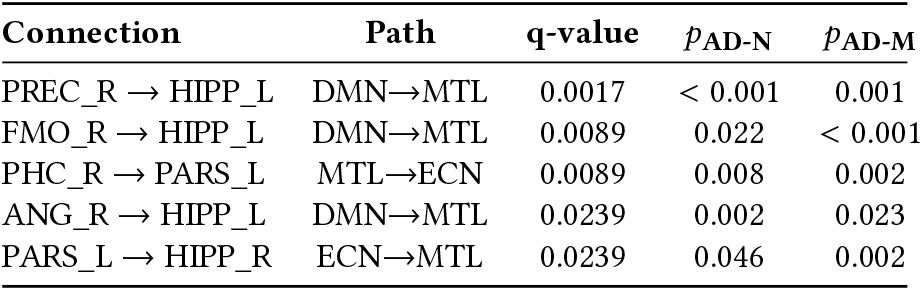
Connections Showing Significant Linear Decrease in Influence Across Disease Stages. Arrows (→) Denote Directed Influence. The Critical Value for Significance is a q-value < 0.05.

**Figure 3:**
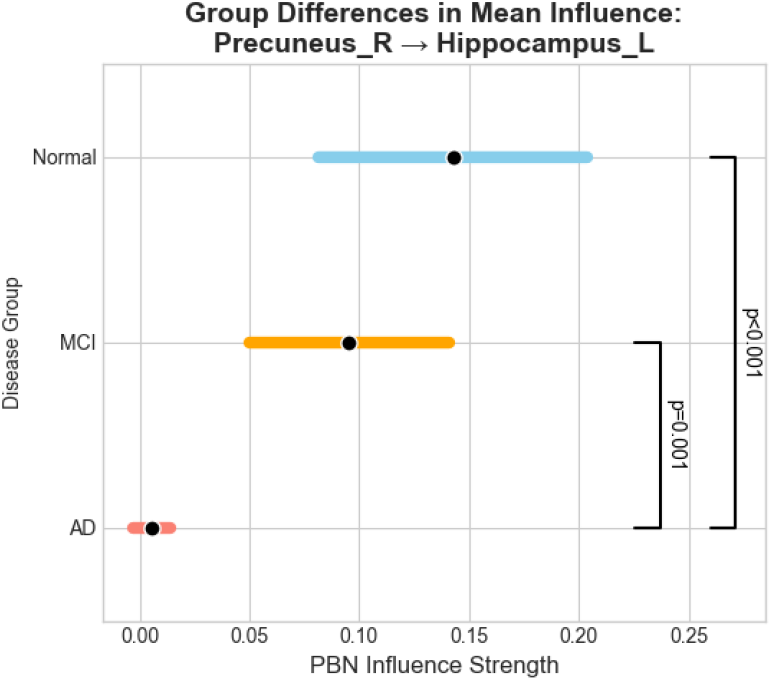
The influence of the Right Precuneus (DMN) on the Left Hippocampus (MTL). The plot shows the mean influence (black circle) and 95% confidence interval for each group. Brackets indicate significant pairwise differences (*p <* 0.05).

**Figure 4:**
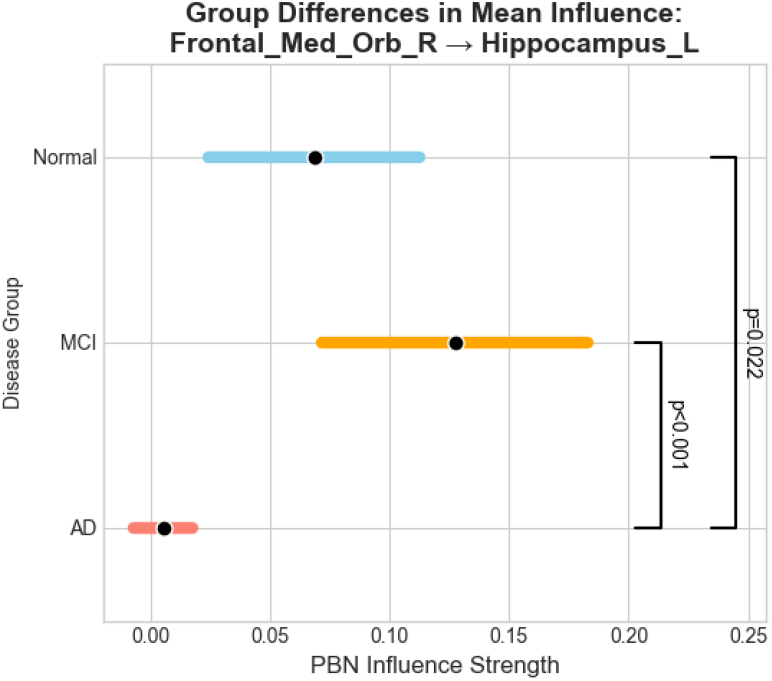
The influence of the Right Frontal Medial Orbital gyrus (DMN) on the Left Hippocampus (MTL). The plot shows the mean influence (black circle) and 95% confidence interval. Brackets indicate significant pairwise differences (*p <* 0.05).

**Figure 5:**
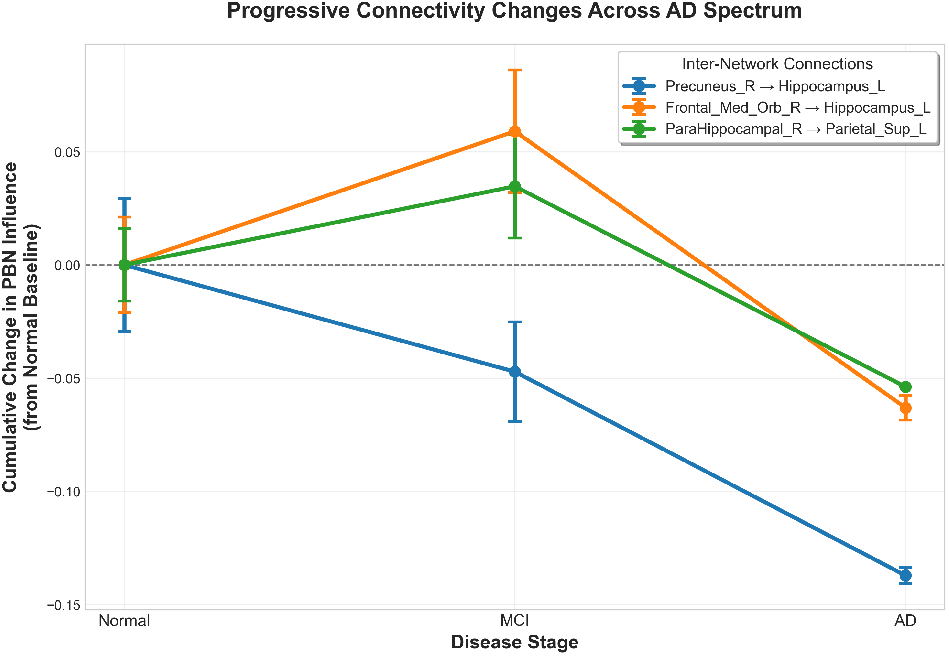
Progressive decline in inter-network connectivity across AD spectrum. Lines show cumulative change in PBN influence strength relative to normal baseline for the three most significant connections (FDR q < 0.05). Error bars represent standard error of the mean. All connections demonstrate systematic weakening from normal controls through MCI to AD, with distinct progression patterns across different network pairs.

**Figure 6:**
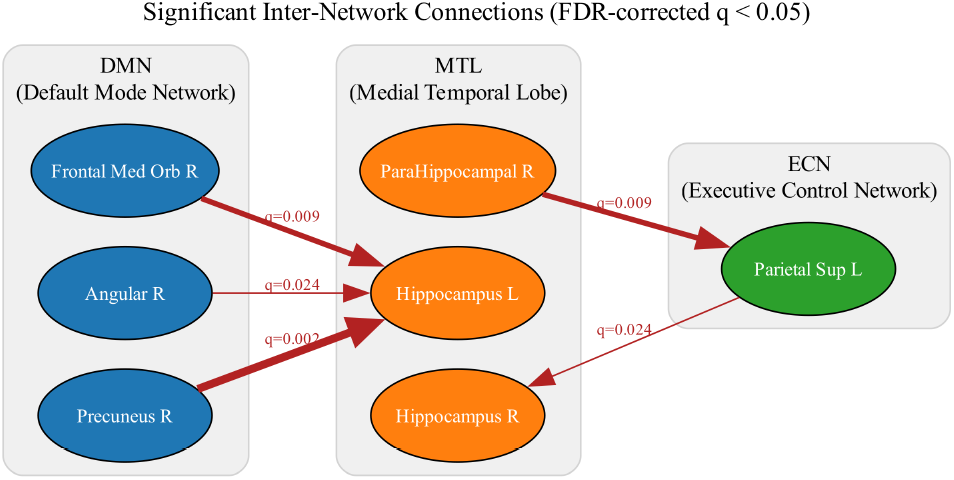
Network diagram of significant inter-network connectivity changes across AD spectrum. Directed edges show connections with significant linear decline in PBN influence (FDR q < 0.05). Edge thickness ∝ − log_10_(q-value). Nodes colored by network: DMN (blue), MTL (orange), ECN (green), SN (red).

The most significant finding was the disrupted influence from the right Precuneus, a key hub of the DMN, to theleft Hippocampus within the MTL (*q* = 0.0017). As shown in Fig. 3 and Fig. 4, the mean influence of this connection is clearly distinct between groups. This pattern of DMN-MTL disconnection is further substantiated by other significant pathways originating from DMN regions, such as the right Medial Orbital Frontal gyrus and the right Angular gyrus, both targeting the left Hippocampus.

### 3.3 Network-Level Interpretation

The overall pattern of results (Fig. 6) points to a cascade of disconnection centered on the brain’s memory system. The systematic failure of three distinct DMN pathways to influence the MTL provides strong, quantifiable evidence for the DMN vulnerability hypothesis in AD [34]. The DMN, a constellation of brain regions including the precuneus and angular gyrus, is central to episodic and autobiographical memory [6]. Critically, this network’s hubs are sites of high metabolic activity, which is thought to underlie their specific vulnerability and their status as primary nodes for early amyloid-*β* deposition, a core pathology of AD [5, 23]. Our findings, which specifically quantify a breakdown in directed influence from DMN nodes like the precuneus to the hippocampus (Table I), therefore model a cornerstone mechanism of AD pathophysiology: the targeted disruption of the primary brain network supporting memory.

The concurrent disruption of ECN-MTL circuits likely contributes to the executive function deficits that accompany memory loss in the disease. The ECN is critical for top-down attentional control, working memory, and goal-directed behavior [29]. While episodic memory impairment is the defining feature of early AD, deficits in executive functions such as planning and cognitive flexibility often manifest as the disease progresses. The observed degradation of pathways between the ECN and MTL (Table I) aligns with models of AD in which the pathology spreads from allocortical (MTL) to association cortices, disrupting the broader networks necessary for higher-order cognition [4]. The failure of ECN-MTL communication thus represents a plausible network-level substrate for the compounding of executive and memory symptoms.

Finally, the consistent linear decrease across all significant findings, clearly visualized for the top connections in Fig. 5, suggests that our PBN framework is effectively capturing the progressive nature of the underlying neurodegenerative process. AD is widely conceptualized not as a static lesion-based disorder but as a progressive disconnection syndrome, where the integrity of large-scale networks degrades over the disease continuum from preclinical, to MCI, and finally to dementia [28]. The fact that our model detected a graded, linear decline in connectivity influence across the NC, MCI, and AD groups for every significant connection provides strong validation. It demonstrates the framework’s sensitivity in capturing the continuous, evolving nature of AD-related neurodegeneration at the network level, moving beyond a simple case-control comparison to map the disease’s progression.

## 4 DISCUSSIONS AND CONCLUSION

We have introduced a novel PBN-based framework for inferring effective brain connectivity that addresses fundamental limitations of existing approaches. Our methodology demonstrates key advantages over both DBNs and DCM: computational tractability enabling large-scale network analysis, preservation of nonlinear logical relationships through Boolean discretization, and robust inference with limited fMRI temporal data. Unlike DCM’s requirement for a priori model specification and DBN’s linear assumptions, our framework successfully extracted meaningful connectivity patterns from standard resting-state sessions while capturing threshold-based neural dynamics. By synergistically integrating data-driven dynamics with anatomical priors, we established a model for progressive disconnection across the AD spectrum.

The key finding was the identification of specific, linear degradation in DMN influence onto the MTL memory system. This provides quantifiable, network-level evidence for a fundamental AD pathophysiological mechanism and demonstrates superior sensitivity compared to correlation-based methods. While DCM excels at hypothesis testing for small, predefined networks, our approach enables data-driven discovery of disease-relevant pathways across selected brain regions without prior model constraints on connectivity patterns. The progressive nature of connectivity loss we observed aligns with established AD neuropathology while providing novel quantitative measures of network dysfunction.

From a clinical perspective, our findings have promising translational implications for AD diagnosis and therapeutic monitoring. The linear connectivity degradation patterns could serve as objective biomarkers for disease staging, potentially enabling earlier detection than current cognitive assessments. These quantitative connectivity measures offer superior sensitivity for tracking subtle network changes that precede overt symptoms, crucial for evaluating therapies in preclinical populations.

For drug development, our methodology provides mechanistic endpoints that could transform clinical trial design. The ability to quantify directed network influences offers objective measures of therapeutic efficacy at the systems level, potentially enabling smaller sample sizes and shorter trial durations. The specific DMNMTL pathway degradation represents a novel therapeutic target, suggesting that interventions aimed at preserving this critical memory circuit could have meaningful clinical benefits.

We acknowledge several limitations that offer avenues for future research. The inherent information loss in BOLD signal binarization represents a simplification of complex neural activity, though this enables computational advantages over continuous modeling approaches. Our individual-structure approach excels at capturing inter-subject variability but may be less powerful for detecting subtle common changes compared to group-level methods. Future studies could explore hybrid models that retain more signal richness while preserving computational efficiency.

Despite these limitations, our structure-aware PBN framework moves beyond correlational analysis to model directed, progressive network failures with greater efficiency and interpretability than existing methods. The specific pathways identified align with established neuropathological models while providing novel targets for understanding disease progression. Our approach establishes PBNs as a powerful tool for translational neuroimaging research with promising clinical relevance.

## ACKNOWLEDGMENTS

The first author would like to express his sincere gratitude to the Undergraduate Research Fellowship Programme (URFP) of the University of Hong Kong for its generous support, which made this research possible. The URFP allowed me to conduct research at the National University of Singapore. The computations were performed using research computing facilities offered by Information Technology Services, the University of Hong Kong.

## A CODE AND DATA AVAILABILITY

All source code used to perform the analyses, along with the meta-data detailing the specific cohorts, are publicly available in a GitHub repository: https://github.com/Liu-Zhonglin/pbn-alzheimers-fmri.

## B COMPLETE ROI MAPPING

**Table 2:** Complete List of 18 Regions of Interest and Network Assignments.

| Abbreviation | AAL3 Region Name | Network |
| --- | --- | --- |
| ANG_L | Angular_L | DMN |
| ANG_R | Angular_R | DMN |
| FMO_L | Frontal_Med_Orb_L | DMN |
| FMO_R | Frontal_Med_Orb_R | DMN |
| PREC_L | Precuneus_L | DMN |
| PREC_R | Precuneus_R | DMN |
| FSG_L | Frontal_Sup_2_L | ECN |
| FSG_R | Frontal_Sup_2_R | ECN |
| PARS_L | Parietal_Sup_L | ECN |
| PARS_R | Parietal_Sup_R | ECN |
| INS_L | Insula_L | SN |
| INS_R | Insula_R | SN |
| SMA_L | Supp_Motor_Area_L | SN |
| SMA_R | Supp_Motor_Area_R | SN |
| HIPP_L | Hippocampus_L | MTL |
| HIPP_R | Hippocampus_R | MTL |
| PHC_L | ParaHippocampal_L | MTL |
| PHC_R | ParaHippocampal_R | MTL |

## Footnotes

1 Data used in the preparation of this article were obtained from the Alzheimer’s Disease Neuroimaging Initiative (ADNI) database (adni.loni.usc.edu). The ADNI was launched in 2003 as a public-private partnership, led by Principal Investigator Michael W. Weiner, MD. The primary goal of ADNI has been to test whether serial magnetic resonance imaging (MRI), positron emission tomography (PET), other biological markers, and clinical and neuropsychological assessments can be combined to measure the progression of MCI and early AD.

