## Supplementary material for "Inferring Progressive Disconnection in Alzheimer’s Disease with Probabilistic Boolean Networks": Tex files: Author-guideline_Wiley.pdf (from another version of this preprint: version 2, 2025-07-18)

### Author guidelines for LaTeX manuscript preparation for Wiley journals in New Journal Design (NJD)

#### 1 Introduction

This document offers step-by-step instructions to prepare LaTeX files using the Wiley NJDv5 LaTeX template. It has been created to help authors prepare LaTeX manuscripts for journals formatted in New Journal Design (NJD), which is Wiley's standard article layout. See the LaTeX page on Wiley's Author Services site where this template is hosted for details about journals using NJD and their font, number of columns, and reference style for authors who would like to simulate these using LaTeX. If the journal to which you want to submit your manuscript is not using NJD, check with the Editorial Office of that journal to ask if they have their own journal-specific LaTeX template. It is possible to use it for journals not using NJD even though it won't look like the final typeset article.

This LaTeX template provides standard coding which Wiley's vendors can successfully convert to XML for typesetting purposes. (Please be aware that manuscript files will be converted by our typesetters into the journal's final specifications for typeset articles, regardless of the reference, font and column number format selected in the LaTeX manuscript.)

This NJDv5 LaTeX template has been created to (1) provide proper guidance to simplify the process, (2) simulate approximately how the article will look once published, and (3) reduce time and manual intervention during the production process which converts the submitted LaTeX manuscript into the journal's final specifications for publication. The template is based on the standard article.cls class file and supports almost all the functionality of that class file.

In the following sections we describe how to install the template package in your system and how to lay out your code using this template to reproduce the typographical look of Wiley NJD journals. If you need support with it, please.

#### 2 Getting started

The WileyNJDv5.cls class file should run on any standard LaTeX installation.

*Installation.* First make sure you have at least MiKTeX 2.9 (21.2) or TeXLive 2021 installed on your computer or use the latest version of your LaTeX editor. Then make sure the WileyNJDv5_template.zip package file is downloaded and extracted to a folder on your computer. The .zip file contains the following:

– WileyNJDv5.cls
– supporting style (.sty) files
– bibliography style (.bst) files based on Wiley reference style
– bibliography (.bib) files based on Wiley reference style sample format

*Usage.* Once this template package is properly installed or copied to the local disk, use the class file WileyNJDv5.cls to create a LaTeX manuscript. Please make sure that your manuscript follows these guidelines.

##### 2.1 Font details

If you would like to use a font that simulates the one used in a specific NJD journal, please go to Wiley's Author Services site where this LaTeX template is hosted and look up the font shown in the spreadsheet for that NJD journal. Alternatively you are free to use the fonts you already have installed in your system. The NJD fonts are as follows:

- Serif fonts – Stix, Times, Garamond, Minion Pro, Utopia, Century, Courier
- Sanserif fonts – Lato, Helvetica, Myriad Pro, Arial, Univers

##### 2.2 How to use the defined fonts

The above listed fonts are available in the fonts folder located within the NJDv5 template folder. Please see screenshot below. There is no need to install any fonts in your system.

**Font folder screenshot**

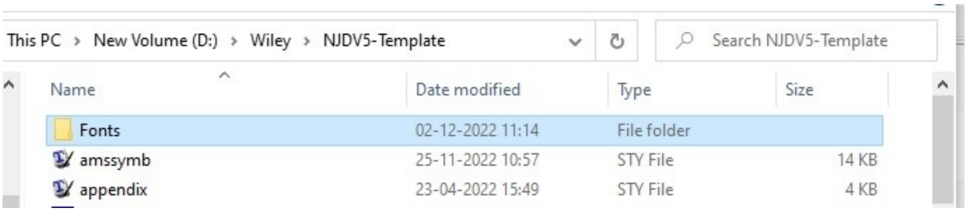


##### 2.3 How to compile the LaTeX file

Based on the LaTeX editor you use, please refer to the respective screenshot given below, and please use the option XeLaTeX for compilation as it supports all NJD fonts. If you are unable to use XeLaTeX, use the default compiler with Times font. On completion of the compilation process, the PDF is available for preview.

**LaTeX editor – TeXworks**

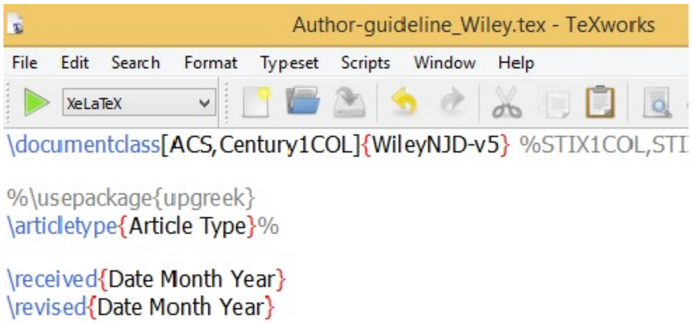

#### LaTeX editor – TeXnicCenter

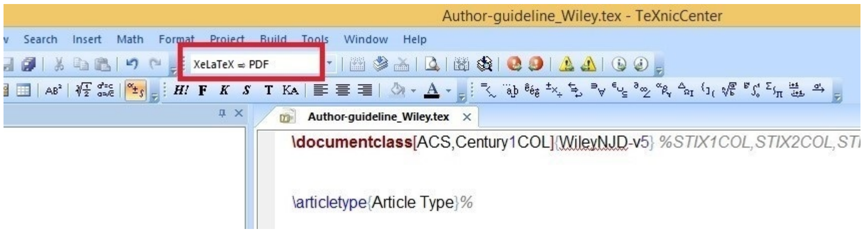


#### LaTeX editor – TeXStudio

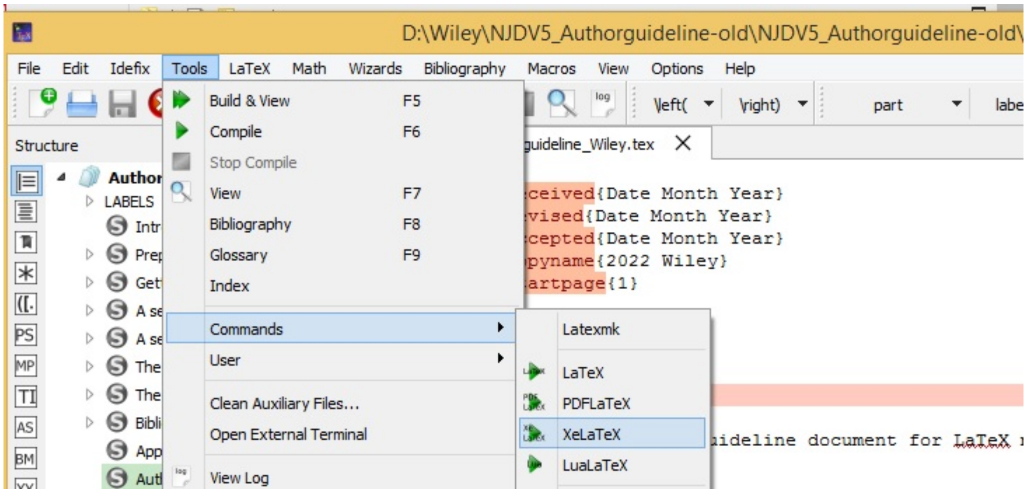


#### LaTeX editor – WinEdt

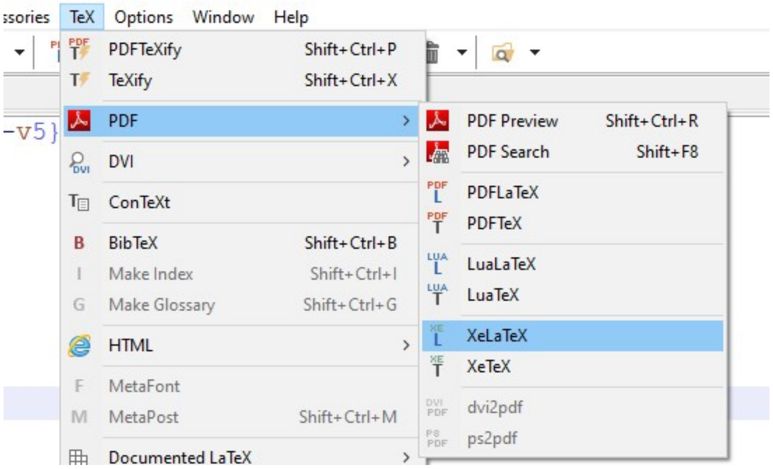

##### 2.4 Usage of the `\documentclass` command

The `\documentclass` command is the first command in the template and defines many options such as one-column format, two-column format, fonts, and reference styles.

You can use a suitable font, reference style, and layout format based on the Wiley journal instructions. For example:

```
\documentclass[AMS,Times1COL]{WileyNJDv5}
- American Mathematical Society reference style
- Times font and 1-column format

\documentclass[AMA,STIX2COL]{WileyNJDv5}
- American Medical Association reference style
- Stix font and 2-column format
```

For further font, layout and reference style options, refer to the Appendix.

#### 3 The Wiley NJDv5 LaTeX template

This template enables you to apply LaTeX coding to your manuscript that will ensure correct formatting of the front matter, body text, and backmatter of the article.

The list of files in the template package includes LaTeX sample, class file, BibTeX style files, supporting files, and font package. The class file WileyNJDv5.cls supports all the NJD styles to simulate the journal layout as closely as possible.

##### 3.1 Preparing your research article

Many researchers prefer to use LaTeX to prepare their manuscript, but this can be difficult to navigate. In these guidelines, we have put together a comprehensive set of LaTeX resources to simplify the process.

Here are some best practice principles to help avoid errors and ensure that the typesetting of your article runs as smoothly as possible. If you are writing for a specific journal, please also check that journal's Author Guidelines on Wiley Online Library for any other journal-level formatting guidelines and instructions.

- Please keep your LaTeX file simple – do not create a complicated preamble containing sophisticated LaTeX constructions. Please keep your own macros to an absolute minimum.
- In particular, please do not change global settings concerning spacings (such as `\parindent`, `\parskip`, `\textwidth`, `\textheight`, and `\pagebreak` etc.) and do not introduce new labels or new environments for definitions, theorems etc.
- As LaTeX is designed to make sensible spacing decisions by itself, please do not use vertical spacing commands, except in a few accepted (mostly mathematical) situations, such as `\bigskip`, `\vskip24pt`, and `\vspace*{24pt}`.
- Please do not use any custom fonts.
- Please make sure that you convert special characters, including diacritical characters such as ä, ö, and ü, into the appropriate LaTeX codes, such as `\''{a}`, `\''{o}`, and `\''{u}`.

More detailed guidance on how to structure all elements of your manuscript using WileyNJDv5.cls is given in Sections 4 and 5.

##### 3.2 Submission of your research article

When it is time to submit your final files, please provide the LaTeX file along with any supporting files used, such as the bibliography/reference (.bib/.bbl) file and the PDF generated from the LaTeX file. Please try to provide your article's references in BibTeX format if possible because this will help during the conversion process if references need to be reformatted to the journal's reference style. Embedded references in the LaTeX file will require the typesetter to carry out a manual process if the references are provided in a format that needs to be converted to the journal's style.

To help with your submission:

- please check for errors in your local compilation before uploading files to the submission system;
- please fix any errors before trying to submit your manuscript, or it may fail to compile.

#### 4 The article header information

An example of article header information using WileyNJDv5.cls is shown below.

```
\documentclass[AMS,STIX1COL]{WileyNJDv5}

\articletype{Article Type}%

\received{00}
\revised{00}
\accepted{00}

\begin{document}

\title{Author guidelines for LaTeX manuscript preparation for
Wiley journals in New Journal Design (NJD)}

\author[1]{Author One}
\author[2,3]{Author Two}
\author[3]{Author Three}

\authormark{Author Name}
\titlemark{Author Guidelines}

\address[1]{\orgdiv{Department Name}, \orgname{Institution Name},
\orgaddress{\state{State Name}, \country{Country Name}}}

\address[2]{\orgdiv{Department Name}, \orgname{Institution Name},
\orgaddress{\state{State Name}, \country{Country Name}}}

\address[3]{\orgdiv{Department Name}, \orgname{Institution Name},
\orgaddress{\state{State Name}, \country{Country Name}}}
```

```
\corres{<Corresponding author information>}

\presentaddress{<author present address information>}

\abstract[Abstract]{This document describes the use of
the LaTeX WileyNJDv5.cls class file for article preparation
for Wiley journals. }

\keywords{Keyword1, Keyword2, Keyword3, Keyword4}

\footnotetext{title footnote...}

\maketitle

\section{Introduction}
.
.
.
```

The compiled output of this article header coding is shown below.

Received: Added at production | Revised: Added at production | Accepted: Added at production

DOI: xxx/xxxx

ARTICLE TYPE

### Author guidelines for LaTeX manuscript preparation for Wiley journals in New Journal Design (NJD)

**Author One[1] | Author Two[2,3] | Author Three[3]**

[1]Department Name, Institution Name, State Name, Country Name

[2]Department Name, Institution Name, State Name, Country Name

[3]Department Name, Institution Name, State Name, Country Name

**Correspondence**
<Corresponding author information>

**Present address**
<author present address information>

**Abstract**

This document describes the use of the LaTeX WileyNJD-v5.cls class file for article preparation for Wiley journals.

KEYWORDS

Keyword1, Keyword2, Keyword3, Keyword4

**Example of an article header information page output**

##### 4.1 Remarks

- Use \authormark{} for running heads.
- Use \received{<received date>} \revised{<revised date>} \accepted{<accepted date>} for history dates. Authors can input these dates as "00" when the manuscript is being prepared.

#### 5 The body of the article

The following sections describe how to code heading levels, equations, tables, figures and other elements which might be needed in the body of the article. Your manuscript should be structured by using the section, subsection and subsubsection environments.

The use of the LaTeX cross-reference system for sections, figures, tables, equations, etc., is encouraged (using \ref{<name>} and \label{<name>}).

##### 5.1 Heading level details

The template is defined by five levels of headings, and the coding for numbered headings is listed below.

Section – use \section{}
Subsection – use \subsection{}
Subsubsection – use \subsubsection{}
Paragraph – use \paragraph{}
Subparagraph – use \subparagraph{}

**4 | A SECTION HEADING**

**4.1 | A subsection heading**

**4.1.1 | A subsubsection heading**

4.1.1.1 | *A paragraph heading*

4.1.1.1.1 | *A subparagraph heading.*

**Examples of numbered section heading levels**

For unnumbered headings, use an asterisk.

Section – use \section*{}
Subsection – use \subsection*{}
Subsubsection – use \subsubsection*{}
Paragraph – use \paragraph*{}
Subparagraph – use \subparagraph*{}

**A SECTION HEADING**

**A subsection heading**

**A subsubsection heading**

*A paragraph heading*

*A subparagraph heading.*

**Examples of unnumbered section heading levels**

#### 5.2 Mathematics: equation coding details

Please ensure that *all* mathematics is correctly coded as maths in the LaTeX file, using e.g. `$...$, $$...$$, \[...\]` or the `equation`, `align`, `eqnarray`, and `gather` environments as appropriate, to enable it to be correctly tagged for online display in the published article. This includes inline maths as well as displayed equations.

##### 5.2.1 Inline equation

Use the standard `$...$` environment to typeset inline equations, for example `$a + b = c.$` to produce the inline equation $a + b = c$.

##### 5.2.2 Display equations

Use the standard `equation` environment to typeset numbered display equations, for example:

```
\begin{equation}
\label{eq1}
a + b = c.
\end{equation}
```

$$a + b = c. \tag{1}$$

Unnumbered centered display equations can be created by using `\[...\]` or `$$...$$` or the `equation*` environment, for example:

```
$${W_{S}} = {\sigma _L}(1 + \cos \theta )
= 2\left( {\sqrt {\sigma _s^d\sigma _L^d}
+ \sqrt {\sigma _s^{nd}\sigma _L^{nd}} } \right)$$
```

$${W_{S}} = {\sigma _L}(1 + \cos \theta ) = 2\left( {\sqrt {\sigma _s^d\sigma _L^d} + \sqrt {\sigma _s^{nd}\sigma _L^{nd}} } \right)$$

For multi-line equations the `align, gather,` or `eqnarray` environment is recommended, for example:

```
\begin{align}
|f(b)-f(a)| &=\left|\int_a^b f'(x)\,{\rm d}x\right|
 \le \int_a^b |f'(x)|\,{\rm d}x\label{eq2}\\
```

```
&\le \int_a^b \max_{a\le t\le b}|f'(t)|\,{\rm d}x
=(b-a)\max_{a\le t\le b}|f'(t)|.
\label{eq3}
\end{align}
```

$$|f(b)-f(a)| = \int_a^b f'(x)\,\mathrm{d}x \le \int_a^b |f'(x)|\,\mathrm{d}x \tag{2}$$

$$\le \int_a^b \max_{a\le t\le b}|f'(t)|\,\mathrm{d}x = (b-a)\max_{a\le t\le b}|f'(t)|. \tag{3}$$

```
\begin{gather}
|f(b)-f(a)| =\left|\int_a^b f'(x)\,{\rm d}x\right|
 \le \int_a^b |f'(x)|\,{\rm d}x\label{eq4}\\
\le \int_a^b \max_{a\le t\le b}|f'(t)|\,{\rm d}x
=(b-a)\max_{a\le t\le b}|f'(t)|.
\label{eq5}
\end{gather}
```

$$|f(b)-f(a)| = \int_a^b f'(x)\,\mathrm{d}x \le \int_a^b |f'(x)|\,\mathrm{d}x \tag{4}$$

$$\le \int_a^b \max_{a\le t\le b}|f'(t)|\,\mathrm{d}x = (b-a)\max_{a\le t\le b}|f'(t)|. \tag{5}$$

```
\begin{eqnarray}
|f(b)-f(a)| &=&\left|\int_a^b f'(x)\,{\rm d}x\right|
 \le \int_a^b |f'(x)|\,{\rm d}x\label{eq6}\\
&\le& \int_a^b \max_{a\le t\le b}|f'(t)|\,{\rm d}x
=(b-a)\max_{a\le t\le b}|f'(t)|.
\label{eq7}
\end{eqnarray}
```

$$|f(b)-f(a)| \;=\; \int_a^b f'(x)\,\mathrm{d}x \le \int_a^b |f'(x)|\,\mathrm{d}x \tag{6}$$

$$\le\; \int_a^b \max_{a\le t\le b}|f'(t)|\,\mathrm{d}x = (b-a)\max_{a\le t\le b}|f'(t)|. \tag{7}$$

For multi-line unnumbered equations the `align*`, `gather*`, or `eqnarray*` environment is recommended, for example:

```
\begin{align*}
|f(b)-f(a)| &=\left|\int_a^b f'(x)\,{\rm d}x\right|
\le \int_a^b |f'(x)|\,{\rm d}x\\
&\le \int_a^b \max_{a\le t\le b}|f'(t)|\,{\rm d}x
=(b-a)\max_{a\le t\le b}|f'(t)|.
\end{align*}
```

$$
\begin{aligned}
|f(b)-f(a)| = \int_a^b f'(x)\,\mathrm{d}x \;\le \int_a^b |f'(x)|\,\mathrm{d}x \\
\le \int_a^b \max_{a\le t\le b}|f'(t)|\,\mathrm{d}x = (b-a)\max_{a\le t\le b}|f'(t)|.
\end{aligned}
$$

```
\begin{gather*}
|f(b)-f(a)| =\left|\int_a^b f'(x)\,{\rm d}x\right|
\le \int_a^b |f'(x)|\,{\rm d}x\\
\le \int_a^b \max_{a\le t\le b}|f'(t)|\,{\rm d}x
=(b-a)\max_{a\le t\le b}|f'(t)|.
\end{gather*}
```

$$
\begin{gathered}
|f(b)-f(a)| = \int_a^b f'(x)\,\mathrm{d}x \;\le \int_a^b |f'(x)|\,\mathrm{d}x \\
\le \int_a^b \max_{a\le t\le b}|f'(t)|\,\mathrm{d}x = (b-a)\max_{a\le t\le b}|f'(t)|.
\end{gathered}
$$

```
\begin{eqnarray*}
|f(b)-f(a)| &=&\left|\int_a^b f'(x)\,{\rm d}x\right|
\le \int_a^b |f'(x)|\,{\rm d}x\\
&\le& \int_a^b \max_{a\le t\le b}|f'(t)|\,{\rm d}x
=(b-a)\max_{a\le t\le b}|f'(t)|.
\end{eqnarray*}
```

$$
\begin{array}{rcl}
|f(b)-f(a)| & = & \int_a^b f'(x)\,\mathrm{d}x \;\le \int_a^b |f'(x)|\,\mathrm{d}x \\
& \le & \int_a^b \max_{a\le t\le b}|f'(t)|\,\mathrm{d}x = (b-a)\max_{a\le t\le b}|f'(t)|.
\end{array}
$$

#### 5.3 Figure and table coding details

WileyNJDv5.cls uses the graphicx/graphics package for handling figures. The standard coding for a figure set in one column is shown below:

```
\begin{figure}
\centering
\includegraphics{filename.eps}
\caption{This is the sample figure caption.}
\label{fig1}
\end{figure}
```

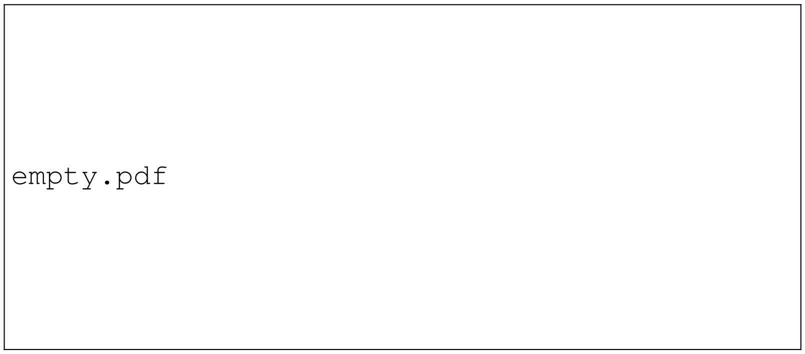


**FIGURE 1** This is the sample figure caption.

Alternatively, if the journal page has two columns, this sample figure coding forces the figure across both columns:

```
\begin{figure*}[!t]
\centerline{\includegraphics{filename.eps}}
\caption{This is the sample figure caption.}
\label{fig2}
\end{figure*}
```

A sample figure citation is: (`Figures~\ref{fig1} and~\ref{fig2}`). When compiled the output will show as Figures 1 and 2.

An example of standard coding for a table is shown below. (Note that the specific example given is for a table being placed on a page using a two-column layout for body text.) The final layout of tables in finalized typeset articles will depend on journal specifications.

```
\begin{table*}%
\caption{This is sample table caption.\label{tab1}}
\begin{tabular*}{\textwidth}{@{\extracolsep\fill}lllll@{}}\toprule
&\multicolumn{2}{@{}c@{}}{\textbf{Spanned heading$^{\tnote{\bf a}}$}}
& \multicolumn{2}{@{}c@{}}{\textbf{Spanned heading$^{\tnote{\bf b}}$}} \\
\cmidrule{2-3}\cmidrule{4-5}
\textbf{Col1 head} & \textbf{Col2 head}  & \textbf{Col3 head}
& \multicolumn{1}{@{}l@{}}{\textbf{Col4 head}}  & \textbf{Col5 head}   \\
\midrule
Col1 text & Col2 text  & Col3 text  & 12.34  & Col5 text\tnote{1}   \\
Col1 text & Col2 text  & Col3 text  & 1.62  & Col5 text\tnote{2}   \\
Col1 text & Col2 text  & Col3 text  & 51.809  & Col5 text   \\
\bottomrule
\end{tabular*}
\begin{tablenotes}
\item[] Example for unnumbered table footnote text.
\item[$^{\rm a}$] Example for a f\kern.01ptirst numbered table footnote.
\item[$^{\rm b}$] Example for a second numbered table footnote.
\end{tablenotes}
\end{table*}
```

**TABLE 1** This is sample table caption.

| | Spanned heading[a] | | Spanned heading[b] | |
|---|---|---|---|---|
| **Col1 head** | **Col2 head** | **Col3 head** | **Col4 head** | **Col5 head** |
| Col1 text | Col2 text | Col3 text | 12.34 | Col5 text1 |
| Col1 text | Col2 text | Col3 text | 1.62 | Col5 text2 |
| Col1 text | Col2 text | Col3 text | 51.809 | Col5 text |

Example for unnumbered table footnote text.
[a]Example for a first numbered table footnote.
[b]Example for a second numbered table footnote.

##### 5.4 Example of coding for display quotes/block quotes

If a display quote or a block quote appears in your manuscript, then use the coding given below.

```
\begin{quote}
This is an example for quote text.

\rightline{---Quote source\hspace*{20pt}}
\end{quote}
```

The output of this coding is shown below:

> This is an example for quote text.
>
> —Quote source

##### 5.5 Examples of boxes with or without a heading

For boxes with or without a heading, the coding details are given below.

```
\begin{boxwithhead}
{BOX 1 This is sample for box heading}
{This is sample for box text. }
\end{boxwithhead}
```

**BOX 1 This is sample for box heading**
This is sample for box text.

```
\begin{boxtext}%
{This is sample for box text. }
\end{boxtext}
```

This is sample for box text.

#### 5.6 List items

##### 5.6.1 Enumerate list styles

```
\begin{enumerate}[1.]
\item list entry
\item list entry
\end{enumerate}

\begin{enumerate}[(1)]
\item list entry
\item list entry
\end{enumerate}

\begin{enumerate}[I.]
\item list entry
\item list entry
\end{enumerate}

\begin{enumerate}[i.]
\item list entry
\item list entry
\end{enumerate}

\begin{enumerate}[(a)]
\item list entry
\item list entry
\end{enumerate}
```

1. list entry
2. list entry

(1) list entry
(2) list entry

I. list entry
II. list entry

i. list entry
ii. list entry

(a) list entry
(b) list entry

##### 5.6.2 Bullet list styles

```
\begin{itemize}
\item bullet list entry
```

```
\item bullet list entry
\end{itemize}
```

- bullet list entry
- bullet list entry

##### 5.6.3 Description list

```
\begin{description}
\item[Step 1] description text.
\item[Step 2] description text.
\end{description}
```

**Step 1** description text.
**Step 2** description text.

#### 5.7 Examples of theorem type environments and proofs

You may use the `claim`, `corollary`, `definition`, `example`, `lemma`, `proposition`, `theorem`, and `remark` environments:

```
\begin{claim}
Claim text goes here.
\end{claim}

\begin{corollary}[Optional Corollary subhead]
\label{cor1}
Corollary text goes here.
\end{corollary}

\begin{definition}[Optional Definition subhead]
\label{def1}
Definition text goes here.
\end{definition}

\begin{example}[Optional Example subhead]
\label{ex1}
Example text goes here.
\end{example}

\begin{lemma}[Optional Lemma subhead]
\label{lem1}
Lemma text goes here.
\end{lemma}

\begin{proposition}[Optional Proposition subhead]
\label{prop1}
```

```
Proposition text goes here.
\end{proposition}

\begin{theorem}[Optional Theorem subhead]
\label{thm1}
Theorem text goes here.
\end{theorem}

\begin{remark}
\label{rem1}
Remark text goes here.
\end{remark}
```

*Claim* 1. Claim text goes here.

**Corollary 1** (Optional Corollary subhead)**.** *Corollary text goes here.*

**Definition 1** (Optional Definition subhead)**.** Definition text goes here.

**Example 1** (Optional Example subhead)**.** Example text goes here.

**Lemma 1** (Optional Lemma subhead)**.** *Lemma text goes here.*

**Proposition 1** (Optional Proposition subhead)**.** *Proposition text goes here.*

**Theorem 1** (Optional Theorem subhead)**.** *Theorem text goes here.*

*Remark* 1. Remark text goes here.

Also available are the environments `assertion`, `conjecture`, `hypothesis`, and `notation`, each of which is numbered by a separate counter:

```
\begin{assertion}
Assertion text goes here.
\end{assertion}

\begin{conjecture}
Conjecture text goes here.
\end{conjecture}

\begin{hypothesis}
Hypothesis text goes here.
\end{hypothesis}

\begin{notation}
Notation text goes here.
\end{notation}
```

**Assertion 1.** *Assertion text goes here.*

**Conjecture 1.** *Conjecture text goes here.*

**Hypothesis 1.** *Hypothesis text goes here.*

*Notation* 1. Notation text goes here.

For proofs please use the `proof` environment.

```
\begin{proof}
Proof text goes here:
\begin{equation*}
 f(b)-f(a)=\int_a^b f'(x)\,{\rm d}x.
\end{equation*}
This completes the proof.
\end{proof}

\begin{proof}[Proof of Theorem~{\rm\ref{thm1}}]
Proof text goes here:
\begin{equation*}
 f(b)-f(a)=\int_a^b f'(x)\,{\rm d}x.
\end{equation*}
This completes the proof.
\end{proof}
```

*Proof.* Proof text goes here:

$$f(b) - f(a) = \int_a^b f'(x)\,\mathrm{d}x.$$

This completes the proof. □

*Proof of Theorem* 1. Proof text goes here:

$$f(b) - f(a) = \int_a^b f'(x)\,\mathrm{d}x.$$

This completes the proof. □

##### 5.8 Program codes

Using the package `listings` you can add non-formatted text as you would do with `\begin{verbatim}` but its main aim is to include the source code of any programming language within your document.
Use `\begin{lstlisting}...\end{lstlisting}` for program codes without mathematics.

The `listings` package supports all the most common languages and it is highly customizable. If you just want to write code within your document, the package provides the `lstlisting` environment; the output will be in Computer Modern typewriter font. Refer to the below example:

```
\begin{lstlisting}[caption={Descriptive caption text},
label=DescriptiveLabel,basicstyle=\fontsize{8}{10}\selectfont\ttfamily]
for i:=maxint to 0 do
begin
{ do nothing }
end;
```

```
Write('Case insensitive ');
Write('Pascal keywords.');
\end{lstlisting}
```

**LISTING 1** Descriptive caption text

```
for i:=maxint to 0 do
begin
{ do nothing }
end;
Write('Case insensitive ');
Write('Pascal keywords.');
```

#### 6 Bibliography: Wiley reference styles

The following .bib format files are provided:

- wileyNJD-WCMS.bib
- wileyNJD-AMA.bib
- wileyNJD-AMS.bib
- wileyNJD-APA.bib
- wileyNJD-APS.bib
- wileyNJD-Chicago.bib
- wileyNJD-Harvard.bib
- wileyNJD-MLA.bib
- wileyNJD-MPS.bib
- wileyNJD-Vancouver.bib

The following bibliography styles are defined:

- wileyNJD-WCMS.bst
- wileyNJD-AMA.bst
- wileyNJD-AMS.bst
- wileyNJD-APA.bst
- wileyNJD-APS.bst
- wileyNJD-Chicago.bst
- wileyNJD-Harvard.bst
- wileyNJD-MLA.bst
- wileyNJD-MPS.bst
- wileyNJD-Vancouver.bst

These 10 BibTeX style (.bst) files are based on the 10 Wiley reference styles: WCMS[1] (Wiley Chemistry–Material Sciences), AMA (American Medical Association), AMS (American Mathematical Society), APA (American

[1]WCMS reference style is based on American Chemical Society (ACS) reference style, which the earlier NJDv2 ACS-Lato and ACS-Stix LaTeX templates included. The Chemistry–Material Sciences reference style isn't a new style, but the same style as used for journals in NJD. ACS reference style has been renamed to WCMS reference style in the NJDv5 template for consistency with the Wiley Journals Style Manual.

Psychological Association), APS (American Physical Society), Chicago, Harvard, MLA (Modern Language Association), MPS (Math and Physical Sciences), and Vancouver.

For example, if AMA reference style is required, select the option AMA in the `\documentclass` command, along with the appropriate font/column option, e.g.

```
\documentclass[AMA,STIX1COL]{WileyNJDv5}
```

(See the LaTeX page on Wiley's Author Services site where this LaTeX template is hosted for details about font, number of columns, and reference style.) Refer to the file wileyNJD-AMA.bib for details on how to provide author name, title, journal name, volume number, issue number, and page number information.

Reference citation commands such as `\cite`, `\citet`, and `\citep` should be used to cross-cite in the body text. Using one of the BibTeX style files provides correct formatting of references as per the Wiley Journals Style Manual. The BibTeX style file identifies the elements of the reference and sets them in the correct style. If a BibTeX style file is not used, and references are included directly in the LaTeX file, when compiled they will be displayed exactly as provided.

Below is an example of how reference citations should be included in the text. This citation coding applies to all reference styles covered by the NJDv5 LaTeX template regardless of whether the journal's citation style is numbered or name-date style.

```
Text with reference citations included \cite{Knupp1999,Kamm2000}.
```

Text with reference citations included (Kamm, 2000; Knupp, 1999).

Examples of the 10 reference styles are given below.

##### 6.1 Example of coding details for Wiley Chemistry–Material Sciences (WCMS) reference style

```
\begin{thebibliography}{10}
\bibitem{Hirt1974} %%%% journal ref.
C. W. Hirt, A. A. Amsden, J. L. Cook, {\it J Comput Phys} \textbf{1974},
{\it 14}, 227.

\bibitem{McWeeny1979} %%%%Book ref.
R. McWeeny, {\it Coulson's Valence}, 3rd ed., Oxford University Press,
Oxford {\bf 1979}.

\bibitem{Schein1992} %%%%Book Series ref.
L. B. Schein, {\it Electrophotography and Development Physics},
2nd ed., Springer Series in Electrophysics, Vol. 14,
Springer, Berlin {\bf 1992}.

\bibitem{Smart1994} %%%%Edited book ref.
A. Smart, in {\it The Chemistry of Metal CVD} (Eds: T. Kodas,
M. Hampden-Smith), VCH, Weinheim, Germany {\bf 1994}, Ch.5.

\bibitem{Author12000} %%%% proceedings
A. B. Author1, C. D. Author2, E. F. Author3, G. H. Author4, in
```

```
{\it Abbrev. Proc. Title} (Eds: I. J. Editor1, K. L. Editor2),
Publisher, Location {\bf Year of publication}, page no.
\end{thebibliography}
```

##### 6.2 Example of coding details for American Medical Association (AMA) reference style

```
\begin{thebibliography}{10}
\bibitem{Hu2002} %%%% journal ref.
Hu P, Reuben DB. Effects of managed care on the length of time that elderly
patients spend with physicians during ambulatory visits: National Ambulatory
Medical Care Survey. {\it Med Care}. 2002;40(7):606-613.
doi:10.1097/00005650-200207000-00007

\bibitem{Geller2002} %%%% Journal with more than 6 authors
Geller AC, Venna S, Prout M, et al. Should the skin cancer examination
be taught in medical school? {\it Arch Dermatol.} 2002;138(9):1201-1203.
doi:10.1001/archderm.138.9.1201

\bibitem{Luketich1995} %%%% book ref.
Luketich JD, Ginsberg RJ. Diagnosis and staging of lung cancer. In: Johnson BE,
Johnson DH, eds. {\it Lung Cancer}. 2nd ed. Wiley-Liss Inc; 1995:161-173.

\bibitem{Slama1994} %%%% Conference/Proceedings ref.
Slama K, ed. Tobacco and Health: Proceedings of the Ninth World Conference on
Tobacco and Health, Paris, France, 10-14 October 1994. Plenum Press; 1995.
\end{thebibliography}
```

##### 6.3 Example of coding details for American Mathematical Society (AMS) reference style

```
\begin{thebibliography}{10}
\bibitem{Li1989} %%% journal ref.
J.-S. Li, {\it Singular unitary representations of classical groups}, Invent.
Math. {\bf 97} (1989), 237--255. MR1001840 (90h:22021).

\bibitem{Harris1996} %%% journal ref. 3 and more than 3 authors
M. Harris, S. S. Kudla, and W. J. Sweet, {\it Theta dichotomy for
unitary groups}, J. Amer. Math. Soc. {\bf 9} (1996),
941--1004. MR1327161 (96m:11041).

\bibitem{Loomis1953} %%%% Book ref.
L. H. Loomis, {\it An introduction to abstract harmonic analysis}, D. Van
Nostrand Company, London, 1953. MR0054173 (14,883c).

\bibitem{Goldman2006} %%% proceeedings ref.
W. M. Goldman, {\it Mapping class group dynamics on surface group
representations}, Problems on mapping class groups and related topics,
Proc. Sympos. Pure Math., vol. 74, Amer. Math. Soc.,
```

```
Providence, RI, 2006, pp. 189--214, DOI 10.1090/pspum/074/2264541. MR2264541.
\end{thebibliography}
```

##### 6.4 Example of coding details for American Psychological Association (APA) reference style

```
\begin{thebibliography}{10}
\bibitem{Grady2019} %%%% Journal ref.
Grady, J. S., Her, M., Moreno, G., Perez, C., \& Yelinek, J. (2019).
Emotions in storybooks: A comparison of storybooks that represent
ethnic and racial groups in the United States. {\it Psychology of
Popular Media Culture}, {\it 8}(3), 207--217.

\bibitem{Jackson2019} %%%% Book ref.
Jackson, L. M. (2019). {\it The psychology of prejudice: From attitudes
to social action} (2nd ed.). American Psychological Association.
https://doi.org/10.1037/0000168-000

\bibitem{Duckworth2019} %%%% proceedings ref.
Duckworth, A. L., Quirk, A., Gallop, R., Hoyle, R. H., Kelly, D. R., \&
Matthews, M. D. (2019). Cognitive and noncognitive predictors of success.
{\it Proceedings of the National Academy of Sciences}, United States,
116(47), 23499--23504. https://doi.org/10.1073/pnas.1910510116
\end{thebibliography}
```

##### 6.5 Example of coding details for American Physical Society (APS) reference style

```
\begin{thebibliography}{10}
\bibitem{Preuss1995} %%%% journal ref.
S. Preuss, A. Demchuk Jr., M. Stuke, Appl. Phys. A {\bf 61}, 33 (1995).

\bibitem{Abrams1973}  %%% book chapter ref.
D. M. Abrams, in {\it Conductive Polymers}, ed. by R. S. Seymour, A. Smith
(Springer, Berlin Heidelberg New York, 1973), p. 307.

\bibitem{Ibach1996} %%% book authored ref.
H. Ibach, H. Lüth, {\it Solid-State Physics}, 2nd ed.
(Springer, New York, 1996).

\bibitem{Zowghi1996} %%%% proceedings ref.
D. Zowghi et al., in {\it PRICAI ’96: Topics in Artificial Intelligence},
ed. by N. Foo, R. Goebel. 4th Pacific Rim Conference on Artificial
Intelligence, Cairns, August 1996. Lecture Notes in Computer Science.
Lecture notes in artificial intelligence, vol. 1114 (Springer, Heidelberg,
1996), p. 157.
\end{thebibliography}
```

##### 6.6 Example of coding details for Chicago reference style

```
\begin{thebibliography}{}
\bibitem{Geoffrey2007} %%%% book ref.
Ward, Geoffrey C., and Ken Burns. 2007. {\it The War: An Intimate History,
1941--1945}. New York: Knopf. Purkis, Samuel, and Victor Klemas. 2011.
{\it Remote Sensing and Global Environmental Change}. Oxford: Wiley-Blackwell.

\bibitem{Weinstein2009} %%%% journal ref.
Weinstein, Joshua I. 2009. ``The Market in Plato's Republic.'' {\it Classical
Philology} 104: 439--58.

\bibitem{Chiswick1977}  %%%%% proceedings ref.
Chiswick, Bake R. 1977. ``A Longitudinal Analysis of the Occupational Mobility
of Immigrants.'' In {\it Proceedings of the 30th Annual Winter Meetings,
Industrial Relations Research Association}, ed. Barbara D. Dennis,
20-7 Madison, WI: IRRA.
\end{thebibliography}
```

##### 6.7 Example of coding details for Harvard reference style

```
\begin{thebibliography}{}
\bibitem{Selman2016}  %%% journal ref.
Selman, P. (2016) The global decline of intercountry adoption:
what lies ahead? {\it Social Policy and Society}, 11(3), 381--397.

\bibitem{Barros2008} %%%% journal ref.
Barros, B., Read, T. \& Verdejo, M.F. (2008) Virtual collaborative
experimentation: an approach combining remote and local labs.
{\it IEEE Transactions on Education},
51(2), 242--250. Available from: https://doi.org/10.1109/TE.2007.908071

\bibitem{Simons2001} %%%% book ref.
Simons, N.E., Menzies, B. \& Matthews, M. (2001) {\it A short course in
soil and rock slope engineering}. London: Thomas Telford Publishing.

\bibitem{Davis2003} %%% book fore more than 7 authors
Davis, M., Charles, S., Curry, M.J., Shanti, H., Prasad, M., Hewings, A.
et al. (2003)  {\it Challenging spatial norms}. London: Routledge.

\bibitem{Wittke2006} %%%% proceedings ref.
Wittke, M. (2006) Design, construction, supervision and long-term behaviour of
tunnels in swelling rock. In: Van Cotthem, A., Charlier, R., Thimus, J.-F. and
Tshibangu, J.-P. (Eds.) Eurock 2006: {\it multiphysics coupling and long term
behaviour in rock mechanics: proceedings of the international symposium of the
international society for rock mechanics, EUROCK 2006,
9-12 May 2006, Liège, Belgium}. London: Taylor \& Francis, pp. 211--216.
\end{thebibliography}
```

##### 6.8 Example of coding details for Modern Language Association (MLA) reference style

```
\begin{thebibliography}{}
\bibitem{Michael1999} %%% book ref.
Dorris, Michael, and Louise Erdrich. {\it The Crown of Columbus}.
HarperCollins Publishers, 1999.

\bibitem{Toorn2017} %%% book ref.
Toorn, Penny van, and Daniel Justice. ``Aboriginal Writing.''
{\it The Cambridge Companion to Canadian Literature},
edited by Eva-Marie Kröller, Cambridge UP, 2017, pp. 26--58.

\bibitem{Kafka2007} %%% journal ref.
Kafka, Ben, and Barbara Adams. ``The Demon of Writing: Paperwork,
Public Safety, and the Reign of Terror.'' {\it Representations},
no. 98, 2007, pp. 1--24.

\bibitem{Chang2000} %%% proceedings ref.
Chang, Steve S., et al., editors. {\it Proceedings of the Twenty-Fifth Annual
Meeting of the Berkeley Linguistics Society, February 12-15, 1999: General
Session and Parasession on Loan Word Phenomena}. Berkeley Linguistics
Society, 2000.
\end{thebibliography}
```

##### 6.9 Example of coding details for Math and Physical Sciences (MPS) reference style

```
\begin{thebibliography}{10}
\bibitem{Hamburger1995} %%%% journal ref.
Hamburger, C.: Quasimonotonicity, regularity and duality for nonlinear
systems of partial differential equations. Ann. Mat. Pura. Appl. 169,
321--354 (1995)

\bibitem{Broy2002} %%%% book ref.
Broy, M.: Software engineering - from auxiliary to key technologies. In:
Broy, M., Denert, E. (eds.) Software Pioneers, pp. 10--13. Springer,
New York (2002)

\bibitem{Zowghi} %%%% proceedings ref.
Zowghi, D., et al.: A framework for reasoning about requirements in evolution.
In: Foo N., Goebel R. (eds.) Topics in Artificial Intelligence,
4th Pacific Rim Conference on Artificial Intelligence, Cairns, August 1996.
Lecture Notes in Computer Science. Lecture Notes in Artificial Intelligence,
vol. 1114, pp. 157--168. Springer, Heidelberg (1996)
\end{thebibliography}
```

##### 6.10 Example of coding details for Vancouver reference style

```
\begin{thebibliography}{10}
\bibitem{Halpern2002}  %%%% journal ref.
Halpern SD, Ubel PA, Caplan AL. Solid-organ transplantation in HIV-infected
patients. N Engl J Med. 2002 Jul 25;347(4):284--7.

\bibitem{Meltzer2002} %%%% book ref.
Meltzer PS, Kallioniemi A, Trent JM. Chromosome alterations in human solid
tumors. In: Vogelstein B, Kinzler KW, editors. The genetic basis of human
cancer. New York: McGraw-Hill; 2002. p. 93--113.

\bibitem{Murray2002} %%%% book ref.
Murray PR, Rosenthal KS, Kobayashi GS, Pfaller MA. Medical microbiology.
4th ed. St. Louis: Mosby; 2002.
\end{thebibliography}
```

#### 7 Appendix

Example of coding details for Appendix headings.

```
\appendix

\bmsection*{Section heading of first appendix\label{app1}}

\bmsubsection*{Subsection heading of f\kern.01ptirst appendix\label{app1.1a}}
```

The output of the above coding is shown below:

**APPENDIX**

**SECTION HEADING OF FIRST APPENDIX**

**Subsection heading of first appendix**

#### `\documentclass` command options for NJD layout styles

Authors can use any one of the reference styles, fonts, and layout format options (given below) in the `\documentclass` command. Please see below layout format with font options.

##### Wiley Chemistry–Material Sciences reference style

```
\documentclass[WCMS,STIX1COL]{WileyNJDv5} % Stix font 1-column format
\documentclass[WCMS,STIX2COL]{WileyNJDv5} % Stix font 2-column format
\documentclass[WCMS,Times1COL]{WileyNJDv5} % Times font 1-column format
\documentclass[WCMS,Times2COL]{WileyNJDv5} % Times font 2-column format
\documentclass[WCMS,Garamond1COL]{WileyNJDv5} % Garamond font 1-column format
\documentclass[WCMS,Garamond2COL]{WileyNJDv5} % Garamond font 2-column format
\documentclass[WCMS,Utopia1COL]{WileyNJDv5} % Utopia font 1-column format
```

```
\documentclass[WCMS,Utopia2COL]{WileyNJDv5} % Utopia font 2-column format
\documentclass[WCMS,Century1COL]{WileyNJDv5} % Century font 1-column format
\documentclass[WCMS,Century2COL]{WileyNJDv5} % Century font 2-column format
\documentclass[WCMS,Minion1COL]{WileyNJDv5} % Minion Pro font 1-column format
\documentclass[WCMS,Minion2COL]{WileyNJDv5} % Minion Pro font 2-column format
\documentclass[WCMS,LATO1COL]{WileyNJDv5} % Lato font 1-column format
\documentclass[WCMS,LATO2COL]{WileyNJDv5} % Lato font 2-column format
\documentclass[WCMS,Helvetica1COL]{WileyNJDv5} % Helvetica font 1-column format
\documentclass[WCMS,Helvetica2COL]{WileyNJDv5} % Helvetica font 2-column format
\documentclass[WCMS,Myriad1COL]{WileyNJDv5} % Myriad font 1-column format
\documentclass[WCMS,Myriad2COL]{WileyNJDv5} % Myriad font 2-column format
\documentclass[WCMS,Arial1COL]{WileyNJDv5} % Arial font 1-column format
\documentclass[WCMS,Arial2COL]{WileyNJDv5} % Arial font 2-column format
\documentclass[WCMS,Univers1COL]{WileyNJDv5} % Univers font 1-column format
\documentclass[WCMS,Univers2COL]{WileyNJDv5} % Univers font 2-column format
\documentclass[WCMS,Courier1COL]{WileyNJDv5} % Courier font 1-column format
\documentclass[WCMS,Courier2COL]{WileyNJDv5} % Courier font 2-column format
```

#### American Mathematical Society reference style

```
\documentclass[AMS,STIX1COL]{WileyNJDv5} % Stix font 1-column format
\documentclass[AMS,STIX2COL]{WileyNJDv5} % Stix font 2-column format
\documentclass[AMS,Times1COL]{WileyNJDv5} % Times font 1-column format
\documentclass[AMS,Times2COL]{WileyNJDv5} % Times font 2-column format
\documentclass[AMS,Garamond1COL]{WileyNJDv5} % Garamond font 1-column format
\documentclass[AMS,Garamond2COL]{WileyNJDv5} % Garamond font 2-column format
\documentclass[AMS,Utopia1COL]{WileyNJDv5} % Utopia font 1-column format
\documentclass[AMS,Utopia2COL]{WileyNJDv5} % Utopia font 2-column format
\documentclass[AMS,Century1COL]{WileyNJDv5} % Century font 1-column format
\documentclass[AMS,Century2COL]{WileyNJDv5} % Century font 2-column format
\documentclass[AMS,Minion1COL]{WileyNJDv5} % Minion Pro font 1-column format
\documentclass[AMS,Minion2COL]{WileyNJDv5} % Minion Pro font 2-column format
\documentclass[AMS,LATO1COL]{WileyNJDv5} % Lato font 1-column format
\documentclass[AMS,LATO2COL]{WileyNJDv5} % Lato font 2-column format
\documentclass[AMS,Helvetica1COL]{WileyNJDv5} % Helvetica font 1-column format
\documentclass[AMS,Helvetica2COL]{WileyNJDv5} % Helvetica font 2-column format
\documentclass[AMS,Myriad1COL]{WileyNJDv5} % Myriad font 1-column format
\documentclass[AMS,Myriad2COL]{WileyNJDv5} % Myriad font 2-column format
\documentclass[AMS,Arial1COL]{WileyNJDv5} % Arial font 1-column format
\documentclass[AMS,Arial2COL]{WileyNJDv5} % Arial font 2-column format
\documentclass[AMS,Univers1COL]{WileyNJDv5} % Univers font 1-column format
\documentclass[AMS,Univers2COL]{WileyNJDv5} % Univers font 2-column format
\documentclass[AMS,Courier1COL]{WileyNJDv5} % Courier font 1-column format
\documentclass[AMS,Courier2COL]{WileyNJDv5} % Courier font 2-column format
```

#### American Medical Association reference style

```
\documentclass[AMA,STIX1COL]{WileyNJDv5} % Stix font 1-column format
\documentclass[AMA,STIX2COL]{WileyNJDv5} % Stix font 2-column format
\documentclass[AMA,Times1COL]{WileyNJDv5} % Times font 1-column format
\documentclass[AMA,Times2COL]{WileyNJDv5} % Times font 2-column format
\documentclass[AMA,Garamond1COL]{WileyNJDv5} % Garamond font 1-column format
\documentclass[AMA,Garamond2COL]{WileyNJDv5} % Garamond font 2-column format
\documentclass[AMA,Utopia1COL]{WileyNJDv5} % Utopia font 1-column format
\documentclass[AMA,Utopia2COL]{WileyNJDv5} % Utopia font 2-column format
\documentclass[AMA,Century1COL]{WileyNJDv5} % Century font 1-column format
\documentclass[AMA,Century2COL]{WileyNJDv5} % Century font 2-column format
\documentclass[AMA,Minion1COL]{WileyNJDv5} % Minion Pro font 1-column format
\documentclass[AMA,Minion2COL]{WileyNJDv5} % Minion Pro font 2-column format
\documentclass[AMA,LATO1COL]{WileyNJDv5} % Lato font 1-column format
\documentclass[AMA,LATO2COL]{WileyNJDv5} % Lato font 2-column format
```

```
\documentclass[AMA,Helvetica1COL]{WileyNJDv5} % Helvetica font 1-column format
\documentclass[AMA,Helvetica2COL]{WileyNJDv5} % Helvetica font 2-column format
\documentclass[AMA,Myriad1COL]{WileyNJDv5} % Myriad font 1-column format
\documentclass[AMA,Myriad2COL]{WileyNJDv5} % Myriad font 2-column format
\documentclass[AMA,Arial1COL]{WileyNJDv5} % Arial font 1-column format
\documentclass[AMA,Arial2COL]{WileyNJDv5} % Arial font 2-column format
\documentclass[AMA,Univers1COL]{WileyNJDv5} % Univers font 1-column format
\documentclass[AMA,Univers2COL]{WileyNJDv5} % Univers font 2-column format
\documentclass[AMA,Courier1COL]{WileyNJDv5} % Courier font 1-column format
\documentclass[AMA,Courier2COL]{WileyNJDv5} % Courier font 2-column format
```

#### American Psychological Association reference style

```
\documentclass[APA,STIX1COL]{WileyNJDv5} % Stix font 1-column format
\documentclass[APA,STIX2COL]{WileyNJDv5} % Stix font 2-column format
\documentclass[APA,Times1COL]{WileyNJDv5} % Times font 1-column format
\documentclass[APA,Times2COL]{WileyNJDv5} % Times font 2-column format
\documentclass[APA,Garamond1COL]{WileyNJDv5} % Garamond font 1-column format
\documentclass[APA,Garamond2COL]{WileyNJDv5} % Garamond font 2-column format
\documentclass[APA,Utopia1COL]{WileyNJDv5} % Utopia font 1-column format
\documentclass[APA,Utopia2COL]{WileyNJDv5} % Utopia font 2-column format
\documentclass[APA,Century1COL]{WileyNJDv5} % Century font 1-column format
\documentclass[APA,Century2COL]{WileyNJDv5} % Century font 2-column format
\documentclass[APA,Minion1COL]{WileyNJDv5} % Minion Pro font 1-column format
\documentclass[APA,Minion2COL]{WileyNJDv5} % Minion Pro font 2-column format
\documentclass[APA,LATO1COL]{WileyNJDv5} % Lato font 1-column format
\documentclass[APA,LATO2COL]{WileyNJDv5} % Lato font 2-column format
\documentclass[APA,Helvetica1COL]{WileyNJDv5} % Helvetica font 1-column format
\documentclass[APA,Helvetica2COL]{WileyNJDv5} % Helvetica font 2-column format
\documentclass[APA,Myriad1COL]{WileyNJDv5} % Myriad font 1-column format
\documentclass[APA,Myriad2COL]{WileyNJDv5} % Myriad font 2-column format
\documentclass[APA,Arial1COL]{WileyNJDv5} % Arial font 1-column format
\documentclass[APA,Arial2COL]{WileyNJDv5} % Arial font 2-column format
\documentclass[APA,Univers1COL]{WileyNJDv5} % Univers font 1-column format
\documentclass[APA,Univers2COL]{WileyNJDv5} % Univers font 2-column format
\documentclass[APA,Courier1COL]{WileyNJDv5} % Courier font 1-column format
\documentclass[APA,Courier2COL]{WileyNJDv5} % Courier font 2-column format
```

#### Vancouver reference style

```
\documentclass[VANCOUVER,STIX1COL]{WileyNJDv5} % Stix font 1-column format
\documentclass[VANCOUVER,STIX2COL]{WileyNJDv5} % Stix font 2-column format
\documentclass[VANCOUVER,Times1COL]{WileyNJDv5} % Times font 1-column format
\documentclass[VANCOUVER,Times2COL]{WileyNJDv5} % Times font 2-column format
\documentclass[VANCOUVER,Garamond1COL]{WileyNJDv5} % Garamond font 1-column format
\documentclass[VANCOUVER,Garamond2COL]{WileyNJDv5} % Garamond font 2-column format
\documentclass[VANCOUVER,Utopia1COL]{WileyNJDv5} % Utopia font 1-column format
\documentclass[VANCOUVER,Utopia2COL]{WileyNJDv5} % Utopia font 2-column format
\documentclass[VANCOUVER,Century1COL]{WileyNJDv5} % Century font 1-column format
\documentclass[VANCOUVER,Century2COL]{WileyNJDv5} % Century font 2-column format
\documentclass[VANCOUVER,Minion1COL]{WileyNJDv5} % Minion Pro font 1-column format
\documentclass[VANCOUVER,Minion2COL]{WileyNJDv5} % Minion Pro font 2-column format
\documentclass[VANCOUVER,LATO1COL]{WileyNJDv5} % Lato font 1-column format
\documentclass[VANCOUVER,LATO2COL]{WileyNJDv5} % Lato font 2-column format
\documentclass[VANCOUVER,Helvetica1COL]{WileyNJDv5} % Helvetica font 1-column format
\documentclass[VANCOUVER,Helvetica2COL]{WileyNJDv5} % Helvetica font 2-column format
\documentclass[VANCOUVER,Myriad1COL]{WileyNJDv5} % Myriad font 1-column format
\documentclass[VANCOUVER,Myriad2COL]{WileyNJDv5} % Myriad font 2-column format
\documentclass[VANCOUVER,Arial1COL]{WileyNJDv5} % Arial font 1-column format
\documentclass[VANCOUVER,Arial2COL]{WileyNJDv5} % Arial font 2-column format
\documentclass[VANCOUVER,Univers1COL]{WileyNJDv5} % Univers font 1-column format
```

```
\documentclass[VANCOUVER,Univers2COL]{WileyNJDv5} % Univers font 2-column format
\documentclass[VANCOUVER,Courier1COL]{WileyNJDv5} % Courier font 1-column format
\documentclass[VANCOUVER,Courier2COL]{WileyNJDv5} % Courier font 2-column format
```

#### Math and Physical Sciences reference style

```
\documentclass[MPS,STIX1COL]{WileyNJDv5} % Stix font 1-column format
\documentclass[MPS,STIX2COL]{WileyNJDv5} % Stix font 2-column format
\documentclass[MPS,Times1COL]{WileyNJDv5} % Times font 1-column format
\documentclass[MPS,Times2COL]{WileyNJDv5} % Times font 2-column format
\documentclass[MPS,Garamond1COL]{WileyNJDv5} % Garamond font 1-column format
\documentclass[MPS,Garamond2COL]{WileyNJDv5} % Garamond font 2-column format
\documentclass[MPS,Utopia1COL]{WileyNJDv5} % Utopia font 1-column format
\documentclass[MPS,Utopia2COL]{WileyNJDv5} % Utopia font 2-column format
\documentclass[MPS,Century1COL]{WileyNJDv5} % Century font 1-column format
\documentclass[MPS,Century2COL]{WileyNJDv5} % Century font 2-column format
\documentclass[MPS,Minion1COL]{WileyNJDv5} % Minion Pro font 1-column format
\documentclass[MPS,Minion2COL]{WileyNJDv5} % Minion Pro font 2-column format
\documentclass[MPS,LATO1COL]{WileyNJDv5} % Lato font 1-column format
\documentclass[MPS,LATO2COL]{WileyNJDv5} % Lato font 2-column format
\documentclass[MPS,Helvetica1COL]{WileyNJDv5} % Helvetica font 1-column format
\documentclass[MPS,Helvetica2COL]{WileyNJDv5} % Helvetica font 2-column format
\documentclass[MPS,Myriad1COL]{WileyNJDv5} % Myriad font 1-column format
\documentclass[MPS,Myriad2COL]{WileyNJDv5} % Myriad font 2-column format
\documentclass[MPS,Arial1COL]{WileyNJDv5} % Arial font 1-column format
\documentclass[MPS,Arial2COL]{WileyNJDv5} % Arial font 2-column format
\documentclass[MPS,Univers1COL]{WileyNJDv5} % Univers font 1-column format
\documentclass[MPS,Univers2COL]{WileyNJDv5} % Univers font 2-column format
\documentclass[MPS,Courier1COL]{WileyNJDv5} % Courier font 1-column format
\documentclass[MPS,Courier2COL]{WileyNJDv5} % Courier font 2-column format
```

#### American Physical Society reference style

```
\documentclass[APS,STIX1COL]{WileyNJDv5} % Stix font 1-column format
\documentclass[APS,STIX2COL]{WileyNJDv5} % Stix font 2-column format
\documentclass[APS,Times1COL]{WileyNJDv5} % Times font 1-column format
\documentclass[APS,Times2COL]{WileyNJDv5} % Times font 2-column format
\documentclass[APS,Garamond1COL]{WileyNJDv5} % Garamond font 1-column format
\documentclass[APS,Garamond2COL]{WileyNJDv5} % Garamond font 2-column format
\documentclass[APS,Utopia1COL]{WileyNJDv5} % Utopia font 1-column format
\documentclass[APS,Utopia2COL]{WileyNJDv5} % Utopia font 2-column format
\documentclass[APS,Century1COL]{WileyNJDv5} % Century font 1-column format
\documentclass[APS,Century2COL]{WileyNJDv5} % Century font 2-column format
\documentclass[APS,Minion1COL]{WileyNJDv5} % Minion Pro font 1-column format
\documentclass[APS,Minion2COL]{WileyNJDv5} % Minion Pro font 2-column format
\documentclass[APS,LATO1COL]{WileyNJDv5} % Lato font 1-column format
\documentclass[APS,LATO2COL]{WileyNJDv5} % Lato font 2-column format
\documentclass[APS,Helvetica1COL]{WileyNJDv5} % Helvetica font 1-column format
\documentclass[APS,Helvetica2COL]{WileyNJDv5} % Helvetica font 2-column format
\documentclass[APS,Myriad1COL]{WileyNJDv5} % Myriad font 1-column format
\documentclass[APS,Myriad2COL]{WileyNJDv5} % Myriad font 2-column format
\documentclass[APS,Arial1COL]{WileyNJDv5} % Arial font 1-column format
\documentclass[APS,Arial2COL]{WileyNJDv5} % Arial font 2-column format
\documentclass[APS,Univers1COL]{WileyNJDv5} % Univers font 1-column format
\documentclass[APS,Univers2COL]{WileyNJDv5} % Univers font 2-column format
\documentclass[APS,Courier1COL]{WileyNJDv5} % Courier font 1-column format
\documentclass[APS,Courier2COL]{WileyNJDv5} % Courier font 2-column format
```

#### Chicago reference style

```
\documentclass[CHICAGO,STIX1COL]{WileyNJDv5} % Stix font 1-column format
\documentclass[CHICAGO,STIX2COL]{WileyNJDv5} % Stix font 2-column format
\documentclass[CHICAGO,Times1COL]{WileyNJDv5} % Times font 1-column format
\documentclass[CHICAGO,Times2COL]{WileyNJDv5} % Times font 2-column format
\documentclass[CHICAGO,Garamond1COL]{WileyNJDv5} % Garamond font 1-column format
\documentclass[CHICAGO,Garamond2COL]{WileyNJDv5} % Garamond font 2-column format
\documentclass[CHICAGO,Utopia1COL]{WileyNJDv5} % Utopia font 1-column format
\documentclass[CHICAGO,Utopia2COL]{WileyNJDv5} % Utopia font 2-column format
\documentclass[CHICAGO,Century1COL]{WileyNJDv5} % Century font 1-column format
\documentclass[CHICAGO,Century2COL]{WileyNJDv5} % Century font 2-column format
\documentclass[CHICAGO,Minion1COL]{WileyNJDv5} % Minion Pro font 1-column format
\documentclass[CHICAGO,Minion2COL]{WileyNJDv5} % Minion Pro font 2-column format
\documentclass[CHICAGO,LATO1COL]{WileyNJDv5} % Lato font 1-column format
\documentclass[CHICAGO,LATO2COL]{WileyNJDv5} % Lato font 2-column format
\documentclass[CHICAGO,Helvetica1COL]{WileyNJDv5} % Helvetica font 1-column format
\documentclass[CHICAGO,Helvetica2COL]{WileyNJDv5} % Helvetica font 2-column format
\documentclass[CHICAGO,Myriad1COL]{WileyNJDv5} % Myriad font 1-column format
\documentclass[CHICAGO,Myriad2COL]{WileyNJDv5} % Myriad font 2-column format
\documentclass[CHICAGO,Arial1COL]{WileyNJDv5} % Arial font 1-column format
\documentclass[CHICAGO,Arial2COL]{WileyNJDv5} % Arial font 2-column format
\documentclass[CHICAGO,Univers1COL]{WileyNJDv5} % Univers font 1-column format
\documentclass[CHICAGO,Univers2COL]{WileyNJDv5} % Univers font 2-column format
\documentclass[CHICAGO,Courier1COL]{WileyNJDv5} % Courier font 1-column format
\documentclass[CHICAGO,Courier2COL]{WileyNJDv5} % Courier font 2-column format
```

#### Harvard reference style

```
\documentclass[HARVARD,STIX1COL]{WileyNJDv5} % Stix font 1-column format
\documentclass[HARVARD,STIX2COL]{WileyNJDv5} % Stix font 2-column format
\documentclass[HARVARD,Times1COL]{WileyNJDv5} % Times font 1-column format
\documentclass[HARVARD,Times2COL]{WileyNJDv5} % Times font 2-column format
\documentclass[HARVARD,Garamond1COL]{WileyNJDv5} % Garamond font 1-column format
\documentclass[HARVARD,Garamond2COL]{WileyNJDv5} % Garamond font 2-column format
\documentclass[HARVARD,Utopia1COL]{WileyNJDv5} % Utopia font 1-column format
\documentclass[HARVARD,Utopia2COL]{WileyNJDv5} % Utopia font 2-column format
\documentclass[HARVARD,Century1COL]{WileyNJDv5} % Century font 1-column format
\documentclass[HARVARD,Century2COL]{WileyNJDv5} % Century font 2-column format
\documentclass[HARVARD,Minion1COL]{WileyNJDv5} % Minion Pro font 1-column format
\documentclass[HARVARD,Minion2COL]{WileyNJDv5} % Minion Pro font 2-column format
\documentclass[HARVARD,LATO1COL]{WileyNJDv5} % Lato font 1-column format
\documentclass[HARVARD,LATO2COL]{WileyNJDv5} % Lato font 2-column format
\documentclass[HARVARD,Helvetica1COL]{WileyNJDv5} % Helvetica font 1-column format
\documentclass[HARVARD,Helvetica2COL]{WileyNJDv5} % Helvetica font 2-column format
\documentclass[HARVARD,Myriad1COL]{WileyNJDv5} % Myriad font 1-column format
\documentclass[HARVARD,Myriad2COL]{WileyNJDv5} % Myriad font 2-column format
\documentclass[HARVARD,Arial1COL]{WileyNJDv5} % Arial font 1-column format
\documentclass[HARVARD,Arial2COL]{WileyNJDv5} % Arial font 2-column format
\documentclass[HARVARD,Univers1COL]{WileyNJDv5} % Univers font 1-column format
\documentclass[HARVARD,Univers2COL]{WileyNJDv5} % Univers font 2-column format
\documentclass[HARVARD,Courier1COL]{WileyNJDv5} % Courier font 1-column format
\documentclass[HARVARD,Courier2COL]{WileyNJDv5} % Courier font 2-column format
```

#### Modern Language Association reference style

```
\documentclass[MLA,STIX1COL]{WileyNJDv5} % Stix font 1-column format
\documentclass[MLA,STIX2COL]{WileyNJDv5} % Stix font 2-column format
\documentclass[MLA,Times1COL]{WileyNJDv5} % Times font 1-column format
\documentclass[MLA,Times2COL]{WileyNJDv5} % Times font 2-column format
\documentclass[MLA,Garamond1COL]{WileyNJDv5} % Garamond font 1-column format
```

```
\documentclass[MLA,Garamond2COL]{WileyNJDv5} % Garamond font 2-column format
\documentclass[MLA,Utopia1COL]{WileyNJDv5} % Utopia font 1-column format
\documentclass[MLA,Utopia2COL]{WileyNJDv5} % Utopia font 2-column format
\documentclass[MLA,Century1COL]{WileyNJDv5} % Century font 1-column format
\documentclass[MLA,Century2COL]{WileyNJDv5} % Century font 2-column format
\documentclass[MLA,Minion1COL]{WileyNJDv5} % Minion Pro font 1-column format
\documentclass[MLA,Minion2COL]{WileyNJDv5} % Minion Pro font 2-column format
\documentclass[MLA,LATO1COL]{WileyNJDv5} % Lato font 1-column format
\documentclass[MLA,LATO2COL]{WileyNJDv5} % Lato font 2-column format
\documentclass[MLA,Helvetica1COL]{WileyNJDv5} % Helvetica font 1-column format
\documentclass[MLA,Helvetica2COL]{WileyNJDv5} % Helvetica font 2-column format
\documentclass[MLA,Myriad1COL]{WileyNJDv5} % Myriad font 1-column format
\documentclass[MLA,Myriad2COL]{WileyNJDv5} % Myriad font 2-column format
\documentclass[MLA,Arial1COL]{WileyNJDv5} % Arial font 1-column format
\documentclass[MLA,Arial2COL]{WileyNJDv5} % Arial font 2-column format
\documentclass[MLA,Univers1COL]{WileyNJDv5} % Univers font 1-column format
\documentclass[MLA,Univers2COL]{WileyNJDv5} % Univers font 2-column format
\documentclass[MLA,Courier1COL]{WileyNJDv5} % Courier font 1-column format
\documentclass[MLA,Courier2COL]{WileyNJDv5} % Courier font 2-column format
```

### 8 Author biography

If the journal requires an author biography, this example below shows how to provide it.

```
\begin{biography}{\includegraphics[width=76pt,height=76pt,draft]{empty}}
{\textbf{Author Name.} Please check with the journal's author guidelines
whether author biographies are required. They are usually only
included for review-type articles, and typically require photos
and brief biographies for each author.}
\end{biography}
```

| | |
|---|---|
| empty.pdf | **Author Name.** Please check with the journal's author guidelines whether author biographies are required. They are usually only included for review-type articles, and typically require photos and brief biographies for each author. |
