## Supplementary material for "Inferring Progressive Disconnection in Alzheimer’s Disease with Probabilistic Boolean Networks": Tex files: wileyNJDv5_AMA.pdf (from another version of this preprint: version 2, 2025-07-18)

Received: Added at production | Revised: Added at production | Accepted: Added at production

DOI: xxx/xxxx

ARTICLE TYPE

### Please insert your article title here

**Mark Taylor**[1] | **Author Two**[2,3] | **Author Three**[3]

[1]Department Name, Institution Name, State Name, Country Name

[2]Department Name, Institution Name, State Name, Country Name

[3]Department Name, Institution Name, State Name, Country Name

**Correspondence**
Corresponding author Mark Taylor, This is sample corresponding address.


**Present address**
This is sample for present address text this is sample for present address text.

**Abstract**

This is a generic template designed for use by multiple journals, which includes several options for customization. Please refer the author guidelines and author LaTeX manuscript preparation document for the journal to which you are submitting in order to confirm that your manuscript will comply with the journal's requirements. Please replace this text with your abstract. This is sample abstract text just for the template display purpose.

**KEYWORDS**
keyword1, keyword2, keyword3, keyword4

#### 1 | FIRST LEVEL HEAD

Please lay out your article using the section headings and the given body text is dummy text for layout purpose. Lorem ipsum dolor sit amet, consectetuer adipiscing elit[1]. Ut purus elit, vestibulum ut, placerat ac, adipiscing vitae, felis. Curabitur dictum gravida mauris. Nam arcu libero, nonummy eget, consectetuer id, vulputate a, magna. Donec vehicula augue eu neque. Pellentesque habitant morbi tristique senectus et netus et malesuada fames ac turpis egestas. Mauris ut leo. Cras viverra metus rhoncus sem. Nulla et lectus vestibulum urna fringilla ultrices. Phasellus eu tellus sit amet tortor gravida placerat. Integer sapien est, iaculis in, pretium quis, viverra ac, nunc. Praesent eget sem vel leo ultrices bibendum. Aenean faucibus. Morbi dolor nulla, malesuada eu, pulvinar at, mollis ac, nulla. Curabitur auctor semper nulla. Donec varius orci eget risus. Duis nibh mi, congue eu, accumsan eleifend, sagittis quis, diam. Duis eget orci sit amet orci dignissim rutrum.

#### 2 | ANOTHER FIRST LEVEL HEAD

Nulla malesuada porttitor diam. Donec felis erat, congue non, volutpat at, tincidunt tristique, libero. Vivamus viverra fermentum felis. Donec nonummy pellentesque ante. Phasellus adipiscing semper elit. Proin fermentum massa ac quam. Sed diam turpis, molestie vitae, placerat a, molestie nec, leo[2]. Maecenas lacinia. Nam ipsum ligula, eleifend at, accumsan nec, suscipit a, ipsum. Morbi blandit ligula feugiat magna. Nunc eleifend consequat lorem. Sed lacinia nulla vitae enim. Pellentesque tincidunt purus vel magna. Integer non enim. Praesent euismod nunc eu purus. Donec bibendum quam in tellus. Nullam cursus pulvinar lectus. Donec et mi. Nam vulputate metus eu enim. Vestibulum pellentesque felis eu massa. Example for bibliography citation[3], text[4,5] inserted in text for your reference.

Quisque ullamcorper placerat ipsum. Cras nibh[6,7]. Morbi vel justo vitae lacus tincidunt ultrices. Lorem ipsum dolor sit amet, consectetuer adipiscing elit. In hac habitasse platea dictumst. Integer tempus convallis augue. Etiam facilisis. Nunc elementum

**Abbreviations:** ANA, anti-nuclear antibodies; APC, antigen-presenting cells; IRF, interferon regulatory factor.

 © 2023 Copyright Holder Name 

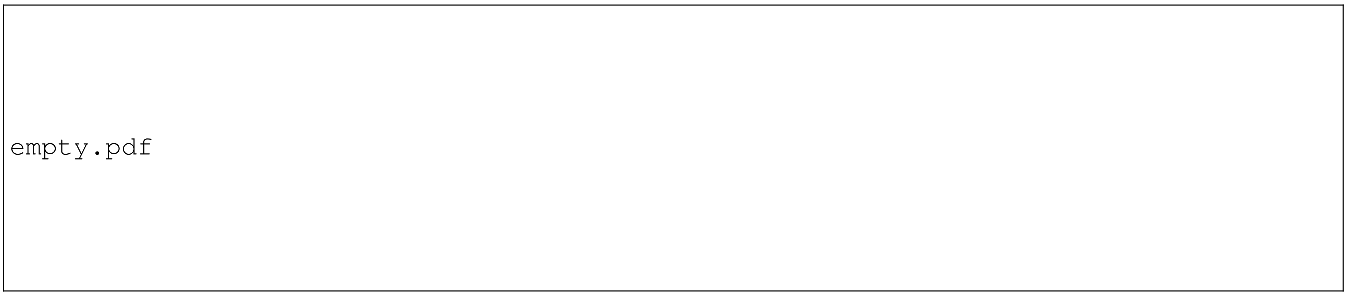


**FIGURE 1** This is the sample figure caption.

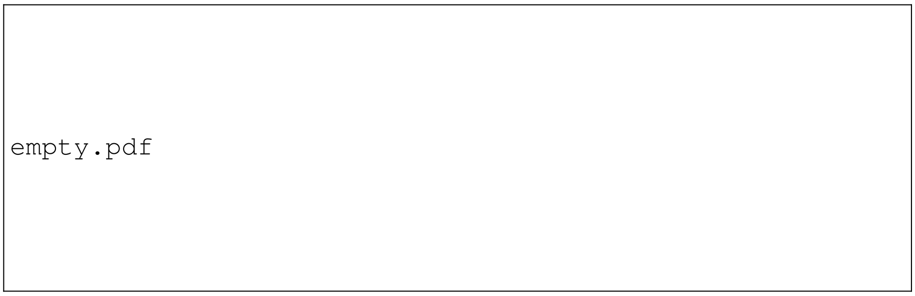


**FIGURE 2** This is the sample figure caption.

fermentum wisi. Aenean placerat. Ut imperdiet, enim sed gravida sollicitudin, felis odio placerat quam, ac pulvinar elit purus eget enim. Nunc vitae tortor. Proin tempus nibh sit amet nisl. Vivamus quis tortor vitae risus porta vehicula.

Fusce mauris. Vestibulum luctus nibh at lectus. Sed bibendum, nulla a faucibus semper, leo velit ultricies tellus, ac venenatis arcu wisi vel nisl. Vestibulum diam. (Figure 1 and 2) Aliquam pellentesque, augue quis sagittis posuere, turpis lacus congues quam, in hendrerit risus eros eget felis. Maecenas eget erat in sapien mattis porttitor. Vestibulum porttitor. Nulla facilisi. Sed a turpis eu lacus commodo facilisis. Morbi fringilla, wisi in dignissim interdum, justo lectus sagittis dui, et vehicula libero dui cursus dui. Mauris tempor ligula sed lacus. Duis cursus enim ut augue. Cras ac magna. Cras nulla. Nulla egestas. Curabitur a leo. Quisque egestas wisi eget nunc. Nam feugiat lacus vel est. Curabitur consectetuer.

Suspendisse vel felis. Ut lorem lorem, interdum eu, tincidunt sit amet, laoreet vitae, arcu. Aenean faucibus pede eu ante. Praesent enim elit, rutrum at, molestie non, nonummy vel, nisl. Ut lectus eros, malesuada sit amet, fermentum eu, sodales cursus, magna. Donec eu purus. Quisque vehicula, urna sed ultricies auctor, pede lorem egestas dui, et convallis elit erat sed nulla. Donec luctus. Curabitur et nunc. Aliquam dolor odio, commodo pretium, ultricies non, pharetra in, velit. Integer arcu est, nonummy in, fermentum faucibus, egestas vel, odio.

$$
\begin{aligned}
s(nT_s) = s(t) \times \sum_{n=0}^{N-1} \delta(t - nT_s) &\xleftrightarrow{\text{DFT}} S\left(\frac{m}{NT_s}\right) \\
&= \frac{1}{N} \sum_{n=0}^{N-1} \sum_{k=-N/2}^{N/2-1} s_k e^{\mathrm{j}2\pi k \Delta f n T_s} e^{-j\frac{2\pi}{N}mn}
\end{aligned}
$$

Sed commodo posuere pede. Mauris ut est. Ut quis purus. Sed ac odio. Sed vehicula hendrerit sem. Duis non odio. Morbi ut dui. Sed accumsan risus eget odio. In hac habitasse platea dictumst. Pellentesque non elit. Fusce sed justo eu urna porta tincidunt. Mauris felis odio, sollicitudin sed, volutpat a, ornare ac, erat. Morbi quis dolor. Donec pellentesque, erat ac sagittis semper, nunc dui lobortis purus, quis congue purus metus ultricies tellus. Proin et quam. Class aptent taciti sociosqu ad litora torquent per conubia nostra, per inceptos hymenaeos. Praesent sapien turpis, fermentum vel, eleifend faucibus, vehicula eu, lacus.

#### 2.1 | Second level head

Pellentesque habitant morbi tristique senectus et netus et malesuada fames ac turpis egestas. Donec odio elit, dictum in, hendrerit sit amet, egestas sed, leo. Praesent feugiat sapien aliquet odio. Integer vitae justo. Aliquam vestibulum fringilla lorem. Sed neque lectus, consectetuer at, consectetuer sed, eleifend ac, lectus. Nulla facilisi. Pellentesque eget lectus. Proin eu metus. Sed porttitor. In hac habitasse platea dictumst. Suspendisse eu lectus. Ut mi mi, lacinia sit amet, placerat et, mollis vitae, dui. Sed ante tellus, tristique ut, iaculis eu, malesuada ac, dui. Mauris nibh leo, facilisis non, adipiscing quis, ultrices a, dui.

Morbi luctus, wisi viverra faucibus pretium, nibh est placerat odio, nec commodo wisi enim eget quam. Quisque libero justo, consectetuer a, feugiat vitae, porttitor eu, libero. Suspendisse sed mauris vitae elit sollicitudin malesuada.

Maecenas ultricies eros sit amet ante. Ut venenatis velit. Maecenas sed mi eget dui varius euismod. Phasellus aliquet volutpat odio. Vestibulum ante ipsum primis in faucibus orci luctus et ultrices posuere cubilia Curae; Pellentesque sit amet pede ac sem eleifend consectetuer. Nullam elementum, urna vel imperdiet sodales, elit ipsum pharetra ligula, ac pretium ante justo a nulla. Curabitur tristique arcu eu metus. Vestibulum lectus. Proin mauris. Proin eu nunc eu urna hendrerit faucibus. Aliquam auctor, pede consequat laoreet varius, eros tellus scelerisque quam, pellentesque hendrerit ipsum dolor sed augue. Nulla nec lacus.

> This is an example[8,9,10] for quote text. This is an example for quote text. This is an example for quote text. This is an example for quote text[11]. This is an example for quote text. This is an example for quote text. This is an example for quote text. This is an example for quote text. This is an example for quote text. This is an example for quote text. This is an example for quote text[12]. This is an example for quote text. This is an example for quote text. This is an example for quote text.

### 3 | EXAMPLE FOR ANOTHER FIRST LEVEL HEAD

#### 3.1 | Example for another second level head

Suspendisse vitae elit. Aliquam arcu neque, ornare in, ullamcorper quis, commodo eu, libero. Fusce sagittis erat at erat tristique mollis. Maecenas sapien libero, molestie et, lobortis in, sodales eget, dui. Morbi ultrices rutrum lorem. Nam elementum ullamcorper leo. Morbi dui. Aliquam sagittis. Nunc placerat. Pellentesque tristique sodales est. Maecenas imperdiet lacinia velit. Cras non urna. Morbi eros pede, suscipit ac, varius vel, egestas non, eros. Praesent malesuada, diam id pretium elementum, eros sem dictum tortor, vel consectetuer odio sem sed wisi.

Sed feugiat. Cum sociis natoque penatibus et magnis dis parturient montes, nascetur ridiculus mus. Ut pellentesque augue sed urna. Vestibulum diam eros, fringilla et, consectetuer eu, nonummy id, sapien. Nullam at lectus. In sagittis ultrices mauris. Curabitur malesuada erat sit amet massa. Fusce blandit. Aliquam erat volutpat. Aliquam euismod. Aenean vel lectus. Nunc imperdiet justo nec dolor.

#### 3.2 | Second level head

Etiam euismod. Fusce facilisis lacinia dui. Suspendisse potenti. In mi erat, cursus id, nonummy sed, ullamcorper eget, sapien. Praesent pretium, magna in eleifend egestas, pede pede pretium lorem, quis consectetuer tortor sapien facilisis magna. Mauris quis magna varius nulla scelerisque imperdiet. Aliquam non quam. Aliquam porttitor quam a lacus. Praesent vel arcu ut tortor cursus volutpat. In vitae pede quis diam bibendum placerat. Fusce elementum convallis neque. Sed dolor orci, scelerisque ac, dapibus nec, ultricies ut, mi. Duis nec dui quis leo sagittis commodo.

##### 3.2.1 | Third level head

Aliquam lectus. Vivamus leo. Quisque ornare tellus ullamcorper nulla. Mauris porttitor pharetra tortor. Sed fringilla justo sed mauris. Mauris tellus. Sed non leo. Nullam elementum, magna in cursus sodales, augue est scelerisque sapien, venenatis congue nulla arcu et pede. Ut suscipit enim vel sapien. Donec congue. Maecenas urna mi, suscipit in, placerat ut, vestibulum ut, massa. Fusce ultrices nulla et nisl.

###### 3.2.1.1 | *Fourth level head*

Sed feugiat. Cum sociis natoque penatibus et magnis dis parturient montes, nascetur ridiculus mus. Ut pellentesque augue sed urna. Vestibulum diam eros, fringilla et, consectetuer eu, nonummy id, sapien. Nullam at lectus. In sagittis ultrices mauris. Curabitur malesuada erat sit amet massa. Fusce blandit. Aliquam erat volutpat. Aliquam euismod. Aenean vel lectus. Nunc imperdiet justo nec dolor.

Etiam euismod. Fusce facilisis lacinia dui. Suspendisse potenti. In mi erat, cursus id, nonummy sed, ullamcorper eget, sapien. Praesent pretium, magna in eleifend egestas, pede pede pretium lorem, quis consectetuer tortor sapien facilisis magna. Mauris quis magna varius nulla scelerisque imperdiet. Aliquam non quam. Aliquam porttitor quam a lacus. Praesent vel arcu ut tortor cursus volutpat. In vitae pede quis diam bibendum placerat. Fusce elementum convallis neque. Sed dolor orci, scelerisque ac, dapibus nec, ultricies ut, mi. Duis nec dui quis leo sagittis commodo.

3.2.1.1.1 | *Fifth level head.* Aliquam lectus. Vivamus leo. Quisque ornare tellus ullamcorper nulla. Mauris porttitor pharetra tortor. Sed fringilla justo sed mauris. Mauris tellus. Sed non leo. Nullam elementum, magna in cursus sodales, augue est scelerisque sapien, venenatis congue nulla arcu et pede. Ut suscipit enim vel sapien. Donec congue. Maecenas urna mi, suscipit in, placerat ut, vestibulum ut, massa. Fusce ultrices nulla et nisl.

Etiam ac leo a risus tristique nonummy. Donec dignissim tincidunt nulla. Vestibulum rhoncus molestie odio. Sed lobortis, justo et pretium lobortis, mauris turpis condimentum augue, nec ultricies nibh arcu pretium enim. Nunc purus neque, placerat id, imperdiet sed, pellentesque nec, nisl. Vestibulum imperdiet neque non sem accumsan laoreet. In hac habitasse platea dictumst. Etiam condimentum facilisis libero. Suspendisse in elit quis nisl aliquam dapibus. Pellentesque auctor sapien. Sed egestas sapien nec lectus. Pellentesque vel dui vel neque bibendum viverra. Aliquam porttitor nisl nec pede. Proin mattis libero vel turpis. Donec rutrum mauris et libero. Proin euismod porta felis. Nam lobortis, metus quis elementum commodo, nunc lectus elementum mauris, eget vulputate ligula tellus eu neque. Vivamus eu dolor.

Nulla in ipsum. Praesent eros nulla, congue vitae, euismod ut, commodo a, wisi. Pellentesque habitant morbi tristique senectus et netus et malesuada fames ac turpis egestas. Aenean nonummy magna non leo. Sed felis erat, ullamcorper in, dictum non, ultricies ut, lectus. Proin vel arcu a odio lobortis euismod. Vestibulum ante ipsum primis in faucibus orci luctus et ultrices posuere cubilia Curae; Proin ut est. Aliquam odio. Pellentesque massa turpis, cursus eu, euismod nec, tempor congue, nulla. Duis viverra gravida mauris. Cras tincidunt. Curabitur eros ligula, varius ut, pulvinar in, cursus faucibus, augue (Box 1).

Etiam ac leo a risus tristique nonummy. Donec dignissim tincidunt nulla. Vestibulum rhoncus molestie odio. Sed lobortis, justo et pretium lobortis, mauris turpis condimentum augue, nec ultricies nibh arcu pretium enim. Nunc purus neque, placerat id, imperdiet sed, pellentesque nec, nisl. Vestibulum imperdiet neque non sem accumsan laoreet. In hac habitasse platea dictumst. Etiam condimentum facilisis libero. Suspendisse in elit quis nisl aliquam dapibus. Pellentesque auctor sapien. Sed egestas sapien nec lectus. Pellentesque vel dui vel neque bibendum viverra. Aliquam porttitor nisl nec pede. Proin mattis libero vel turpis. Donec rutrum mauris et libero. Proin euismod porta felis. Nam lobortis, metus quis elementum commodo, nunc lectus elementum mauris, eget vulputate ligula tellus eu neque. Vivamus eu dolor.

**BOX 1 This is sample for Box head and text**

This is sample for boxtext this is sample for boxtext this is sample for boxtext this is sample for boxtext this is sample for boxtext this is sample for boxtext this is sample for boxtext this is sample for boxtext this is sample for boxtext this is sample for boxtext this is sample for boxtext this is sample for boxtext this is sample for boxtext this is sample for boxtext this is sample for boxtext this is sample for boxtext this is sample for boxtext this is sample for boxtext this is sample for boxtext this is sample for boxtext this is sample for boxtext this is sample for boxtext this is sample for boxtext this is sample for boxtext this is sample for boxtext this is sample for boxtext this is sample for boxtext this is sample for boxtext this is sample for boxtext this is sample for boxtext this is sample for boxtext this is sample for boxtext this is sample for boxtext this is sample for boxtext this is sample for boxtext this is sample for boxtext this is sample for boxtext this is sample for boxtext this is sample for boxtext this is sample for boxtext this is sample for boxtext this is sample for boxtext this is sample for boxtext this is sample for boxtext this is sample for boxtext this is sample for boxtext this is sample for boxtext this is sample for boxtext this is sample for boxtext this is sample for boxtext this is sample for boxtext this is sample for boxtext. This is sample for boxtext this is sample for boxtext this is sample for boxtext this is sample for boxtext this is sample for boxtext this is sample for boxtext this is sample for boxtext this is sample for boxtext this is sample for boxtext this is sample for boxtext this is sample for boxtext this is sample for boxtext this is sample for boxtext this is sample for boxtext this is sample for boxtext this is sample for boxtext this is sample for boxtext this is sample for boxtext this is sample for boxtext this is sample for boxtext this is sample for boxtext this is sample for boxtext this is sample for boxtext this is sample for boxtext this is sample for boxtext this is sample for boxtext this is sample for boxtext this is sample for boxtext this is sample for boxtext this is sample for boxtext this is sample for boxtext this is sample for boxtext this is sample for boxtext this is sample for boxtext this is sample for boxtext this is sample for boxtext this is sample for boxtext this is sample for boxtext this is sample for boxtext this is sample for boxtext this is sample for boxtext this is sample for boxtext this is sample for boxtext this is sample for boxtext this is sample for boxtext this is sample for boxtext this is sample for boxtext this is sample for boxtext this is sample for boxtext this is sample for boxtext this is sample for boxtext.

**TABLE 1** This is sample table caption.

| | Spanned heading[a] | | Spanned heading[b] | |
|---|---|---|---|---|
| **Col1 head** | **Col2 head** | **Col3 head** | **Col4 head** | **Col5 head** |
| col1 text | col2 text | col3 text | 12.34 | col5 text1 |
| col1 text | col2 text | col3 text | 1.62 | col5 text2 |
| col1 text | col2 text | col3 text | 51.809 | col5 text |

[a]Example for a first table footnote.
[b]Example for a second table footnote.
*Source*: Example for table source text.

Curabitur tellus magna, porttitor a, commodo a, commodo in, tortor. Donec interdum (Table 1). Praesent scelerisque. Maecenas posuere sodales odio. Vivamus metus lacus, varius quis, imperdiet quis, rhoncus a, turpis. Etiam ligula arcu, elementum a, venenatis quis, sollicitudin sed, metus. Donec nunc pede, tincidunt in, venenatis vitae, faucibus vel, nibh. Pellentesque wisi. Nullam malesuada. Morbi ut tellus ut pede tincidunt porta. Lorem ipsum dolor sit amet, consectetuer adipiscing elit. Etiam congue neque id dolor.

Donec et nisl at wisi luctus bibendum. Nam interdum tellus ac libero. Sed sem justo, laoreet vitae, fringilla at, adipiscing ut, nibh. Maecenas non sem quis tortor eleifend fermentum. Etiam id tortor ac mauris porta vulputate. Integer porta neque vitae massa. Maecenas tempus libero a libero posuere dictum. Vestibulum ante ipsum primis in faucibus orci luctus et ultrices posuere cubilia Curae; Aenean quis mauris sed elit commodo placerat. Class aptent taciti sociosqu ad litora torquent per conubia nostra, per inceptos hymenaeos. Vivamus rhoncus tincidunt libero. Etiam elementum pretium justo. Vivamus est. Morbi a tellus eget pede tristique commodo. Nulla nisl. Vestibulum sed nisl eu sapien cursus rutrum.

This is sample for boxtext this is sample for boxtext this is sample for boxtext this is sample for boxtext this is sample for boxtext this is sample for boxtext this is sample for boxtext this is sample for boxtext this is sample for boxtext this is sample for boxtext this is sample for boxtext this is sample for boxtext this is sample for boxtext this is sample for boxtext this is sample for boxtext this is sample for boxtext.

Nulla non mauris vitae wisi posuere convallis. Sed eu nulla nec eros scelerisque pharetra. Nullam varius. Etiam dignissim elementum metus. Vestibulum faucibus, metus sit amet mattis rhoncus, sapien dui laoreet odio, nec ultricies nibh augue a enim. Fusce in ligula. Quisque at magna et nulla commodo consequat. Proin accumsan imperdiet sem. Nunc porta. Donec feugiat mi at justo. Phasellus facilisis ipsum quis ante. In ac elit eget ipsum pharetra faucibus. Maecenas viverra nulla in massa (Table 2).

Nulla in ipsum. Praesent eros nulla, congue vitae, euismod ut, commodo a, wisi. Pellentesque habitant morbi tristique senectus et netus et malesuada fames ac turpis egestas. Aenean nonummy magna non leo. Sed felis erat, ullamcorper in, dictum non, ultricies ut, lectus. Proin vel arcu a odio lobortis euismod. Vestibulum ante ipsum primis

Fusce mauris. Vestibulum luctus nibh at lectus. Sed bibendum, nulla a faucibus semper, leo velit ultricies tellus, ac venenatis arcu wisi vel nisl. Vestibulum diam. Aliquam pellentesque, augue quis sagittis posuere, turpis lacus congue quam, in hendrerit risus eros eget felis. Maecenas eget erat in sapien mattis porttitor. Vestibulum porttitor. Nulla facilisi. Sed a turpis eu lacus commodo facilisis. Morbi fringilla, wisi in dignissim interdum, justo lectus sagittis dui, et vehicula libero dui cursus dui. Mauris

**TABLE 2** This is sample table caption.

| **Col1 head** | **Col2 head** | **Col3 head** | **Col4 head** | **Col5 head** |
|---|---|---|---|---|
| col1 text | col2 text | col3 text | col4 text | col5 text† |
| col1 text | col2 text | col3 text | col4 text | col5 text |
| col1 text | col2 text | col3 text | col4 text | col5 text‡ |

†Example for a first table footnote.
‡Example for a second table footnote.
*Source*: Example for table source text.

tempor ligula sed lacus. Duis cursus enim ut augue. Cras ac magna. Cras nulla. Nulla egestas. Curabitur a leo. Quisque egestas wisi eget nunc. Nam feugiat lacus vel est. Curabitur consectetuer.

Below is the example[2,6,7] for bulleted list. Below is the example for bulleted list. Below is the example for bulleted list. Below is the example for bulleted list. Below is the example for bulleted list. Below is the example for bulleted list‡:

- bulleted list entry sample bulleted list entry[13], sample list entry text.
- bulleted list entry sample bulleted list entry. bulleted list entry sample bulleted list entry. bulleted list entry sample bulleted list entry.
- bulleted list entry sample bulleted list entry[14], bulleted list entry sample bulleted list entry[15], sample list entry text. bulleted list entry sample bulleted list entry.
- sample list entry text. sample list entry text.

Suspendisse vel felis. Ut lorem lorem, interdum eu, tincidunt sit amet, laoreet vitae, arcu. Aenean faucibus pede eu ante. Praesent enim elit, rutrum at, molestie non, nonummy vel, nisl. Ut lectus eros, malesuada sit amet, fermentum eu, sodales cursus, magna. Donec eu purus. Quisque vehicula, urna sed ultricies auctor, pede lorem egestas dui, et convallis elit erat sed nulla. Donec luctus. Curabitur et nunc. Aliquam dolor odio, commodo pretium, ultricies non, pharetra in, velit. Integer arcu est, nonummy in, fermentum faucibus, egestas vel, odio.

Sed commodo posuere pede. Mauris ut est. Ut quis purus. Sed ac odio. Sed vehicula hendrerit sem. Duis non odio. Morbi ut dui. Sed accumsan risus eget odio. In hac habitasse platea dictumst. Pellentesque non elit. Fusce sed justo eu urna porta tincidunt. Mauris felis odio, sollicitudin sed, volutpat a, ornare ac, erat. Morbi quis dolor. Donec pellentesque, erat ac sagittis semper, nunc dui lobortis purus, quis congue purus metus ultricies tellus. Proin et quam. Class aptent taciti sociosqu ad litora torquent per conubia nostra, per inceptos hymenaeos. Praesent sapien turpis, fermentum vel, eleifend faucibus, vehicula eu, lacus.

Pellentesque non elit. Fusce sed justo eu urna porta tincidunt. Mauris felis odio, sollicitudin sed, volutpat a, ornare ac, erat. Morbi quis dolor. Donec pellentesque, erat ac sagittis semper, nunc dui lobortis purus, quis congue purus metus ultricies tellus. Proin et quam. Below is the example for description list. Below is the example for description list. Below is the example for description list. Below is the example for description list. Below is the example for description list. Below is the sample for description list. Below is the example for description list. Below is the example for description list. Below is the example for description list. Below is the example for description list. Below is the example for description list:

**Description sample:**

**first entry** description text, description text[16,17,18]. description text, description text, description text, description text, description text.
**second long entry** description text, description text, description text, description text, description text, description text, description text.
**third entry** description text, description text, description text, description text, description text.
**fourth entry** description text, description text.

**Numbered list items sample:**

1. First level numbered list entry, sample numbered list entry.

‡ This is an example for footnote.

2. First numbered list entry, sample numbered list entry. Numbered list entry [19], sample numbered list entry. Numbered list entry, sample numbered list entry.
   a. Second level alpabetical list entry. Second level alpabetical list entry. Second level alpabetical list entry [20]. Second level alpabetical list entry.
   b. Second level alpabetical list entry. Second level alpabetical list entry [21,4,4].
      i. Third level lowercase roman numeral list entry. Third level lowercase roman numeral list entry. Third level lowercase roman numeral list entry.
      ii. Third level lowercase roman numeral list entry. Third level lowercase roman numeral list entry. [5]
   c. Second level alpabetical list entry. Second level alpabetical list entry [3].
3. First level numbered list entry, sample numbered list entry. Numbered list entry, sample numbered list entry. Numbered list entry.
4. Another first level numbered list entry, sample numbered list entry. Numbered list entry, sample numbered list entry. Numbered list entry.

**un-numbered list items sample:**

Sample unnumberd list text.
Sample unnumberd list text.
sample unnumberd list text.
Sample unnumberd list text.

#### 4 | EXAMPLES FOR ENUNCIATIONS

**Theorem 1.** *Example theorem text. Example theorem text. Example theorem text. Example theorem text. Example theorem text. Example theorem text. Example theorem text. Example theorem text. Example theorem text. Example theorem text. Example theorem text. Example theorem text. Example theorem text. Example theorem text. Example theorem text. Example theorem text. Example theorem text. Example theorem text. Example theorem text. Example theorem text. Example theorem text. Example theorem text. Example theorem text. Example theorem text. Example theorem text.*

Quisque ullamcorper placerat ipsum. Cras nibh. Morbi vel justo vitae lacus tincidunt ultrices. Lorem ipsum dolor sit amet, consectetuer adipiscing elit. In hac habitasse platea dictumst. Integer tempus convallis augue. Etiam facilisis. Nunc elementum fermentum wisi. Aenean placerat. Ut imperdiet, enim sed gravida sollicitudin, felis odio placerat quam, ac pulvinar elit purus eget enim. Nunc vitae tortor. Proin tempus nibh sit amet nisl. Vivamus quis tortor vitae risus porta vehicula.

Fusce mauris. Vestibulum luctus nibh at lectus. Sed bibendum, nulla a faucibus semper, leo velit ultricies tellus, ac venenatis arcu wisi vel nisl. Vestibulum diam. Aliquam pellentesque, augue quis sagittis posuere, turpis lacus congue quam, in hendrerit risus eros eget felis. Maecenas eget erat in sapien mattis porttitor. Vestibulum porttitor. Nulla facilisi. Sed a turpis eu lacus commodo facilisis. Morbi fringilla, wisi in dignissim interdum, justo lectus sagittis dui, et vehicula libero dui cursus dui. Mauris tempor ligula sed lacus. Duis cursus enim ut augue. Cras ac magna. Cras nulla. Nulla egestas. Curabitur a leo. Quisque egestas wisi eget nunc. Nam feugiat lacus vel est. Curabitur consectetuer.

**Proposition 1.** *Example proposition text. Example proposition text. Example proposition text. Example proposition text. Example proposition text. Example proposition text. Example proposition text. Example proposition text. Example proposition text. Example proposition text. Example proposition text. Example proposition text. Example proposition text. Example proposition text. Example proposition text. Example proposition text.*

Nulla malesuada porttitor diam. Donec felis erat, congue non, volutpat at, tincidunt tristique, libero. Vivamus viverra fermentum felis. Donec nonummy pellentesque ante. Phasellus adipiscing semper elit. Proin fermentum massa ac quam. Sed diam turpis, molestie vitae, placerat a, molestie nec, leo. Maecenas lacinia. Nam ipsum ligula, eleifend at, accumsan nec, suscipit a, ipsum. Morbi blandit ligula feugiat magna. Nunc eleifend consequat lorem. Sed lacinia nulla vitae enim. Pellentesque tincidunt purus vel magna. Integer non enim. Praesent euismod nunc eu purus. Donec bibendum quam in tellus. Nullam cursus pulvinar lectus. Donec et mi. Nam vulputate metus eu enim. Vestibulum pellentesque felis eu massa.

Quisque ullamcorper placerat ipsum. Cras nibh. Morbi vel justo vitae lacus tincidunt ultrices. Lorem ipsum dolor sit amet, consectetuer adipiscing elit. In hac habitasse platea dictumst. Integer tempus convallis augue. Etiam facilisis. Nunc elementum fermentum wisi. Aenean placerat. Ut imperdiet, enim sed gravida sollicitudin, felis odio placerat quam, ac pulvinar elit purus eget enim. Nunc vitae tortor. Proin tempus nibh sit amet nisl. Vivamus quis tortor vitae risus porta vehicula.

**Definition 1.** Example definition text. Example definition text. Example definition text. Example definition text. Example definition text. Example definition text. Example definition text. Example definition text. Example definition text. Example definition text. Example definition text.

Sed commodo posuere pede. Mauris ut est. Ut quis purus. Sed ac odio. Sed vehicula hendrerit sem. Duis non odio. Morbi ut dui. Sed accumsan risus eget odio. In hac habitasse platea dictumst. Pellentesque non elit. Fusce sed justo eu urna porta tincidunt. Mauris felis odio, sollicitudin sed, volutpat a, ornare ac, erat. Morbi quis dolor. Donec pellentesque, erat ac sagittis semper, nunc dui lobortis purus, quis congue purus metus ultricies tellus. Proin et quam. Class aptent taciti sociosqu ad litora torquent per conubia nostra, per inceptos hymenaeos. Praesent sapien turpis, fermentum vel, eleifend faucibus, vehicula eu, lacus.

Pellentesque habitant morbi tristique senectus et netus et malesuada fames ac turpis egestas. Donec odio elit, dictum in, hendrerit sit amet, egestas sed, leo. Praesent feugiat sapien aliquet odio. Integer vitae justo. Aliquam vestibulum fringilla lorem. Sed neque lectus, consectetuer at, consectetuer sed, eleifend ac, lectus. Nulla facilisi. Pellentesque eget lectus. Proin eu metus. Sed porttitor. In hac habitasse platea dictumst. Suspendisse eu lectus. Ut mi mi, lacinia sit amet, placerat et, mollis vitae, dui. Sed ante tellus, tristique ut, iaculis eu, malesuada ac, dui. Mauris nibh leo, facilisis non, adipiscing quis, ultrices a, dui.

*Proof.* Example for proof text. Example for proof text. Example for proof text. Example for proof text. Example for proof text. Example for proof text. Example for proof text. Example for proof text. Example for proof text. Example for proof text. □

Nam dui ligula, fringilla a, euismod sodales, sollicitudin vel, wisi. Morbi auctor lorem non justo. Nam lacus libero, pretium at, lobortis vitae, ultricies et, tellus. Donec aliquet, tortor sed accumsan bibendum, erat ligula aliquet magna, vitae ornare odio metus a mi. Morbi ac orci et nisl hendrerit mollis. Suspendisse ut massa. Cras nec ante. Pellentesque a nulla. Cum sociis natoque penatibus et magnis dis parturient montes, nascetur ridiculus mus. Aliquam tincidunt urna. Nulla ullamcorper vestibulum turpis. Pellentesque cursus luctus mauris.

Nulla malesuada porttitor diam. Donec felis erat, congue non, volutpat at, tincidunt tristique, libero. Vivamus viverra fermentum felis. Donec nonummy pellentesque ante. Phasellus adipiscing semper elit. Proin fermentum massa ac quam. Sed diam turpis, molestie vitae, placerat a, molestie nec, leo. Maecenas lacinia. Nam ipsum ligula, eleifend at, accumsan nec, suscipit a, ipsum. Morbi blandit ligula feugiat magna. Nunc eleifend consequat lorem. Sed lacinia nulla vitae enim. Pellentesque tincidunt purus vel magna. Integer non enim. Praesent euismod nunc eu purus. Donec bibendum quam in tellus. Nullam cursus pulvinar lectus. Donec et mi. Nam vulputate metus eu enim. Vestibulum pellentesque felis eu massa.

*Proof of Theorem* 1. Example for proof text. Example for proof text. Example for proof text. Example for proof text. Example for proof text. Example for proof text. Example for proof text. Example for proof text. Example for proof text. Example for proof text. □

Etiam euismod. Fusce facilisis lacinia dui. Suspendisse potenti. In mi erat, cursus id, nonummy sed, ullamcorper eget, sapien. Praesent pretium, magna in eleifend egestas, pede pede pretium lorem, quis consectetuer tortor sapien facilisis magna. Mauris quis magna varius nulla scelerisque imperdiet. Aliquam non quam. Aliquam porttitor quam a lacus. Praesent vel arcu ut tortor cursus volutpat. In vitae pede quis diam bibendum placerat. Fusce elementum convallis neque. Sed dolor orci, scelerisque ac, dapibus nec, ultricies ut, mi. Duis nec dui quis leo sagittis commodo. Aliquam lectus. Vivamus leo. Quisque ornare tellus ullamcorper nulla. Mauris porttitor pharetra tortor. Sed fringilla justo sed mauris. Mauris tellus. Sed non leo. Nullam elementum, magna in cursus sodales, augue est scelerisque sapien, venenatis congue nulla arcu et pede. Ut suscipit enim vel sapien. Donec congue. Maecenas urna mi, suscipit in, placerat ut, vestibulum ut, massa. Fusce ultrices nulla et nisl (Table 3).

Pellentesque habitant morbi tristique senectus et netus et malesuada fames ac turpis egestas. Donec odio elit, dictum in, hendrerit sit amet, egestas sed, leo. Praesent feugiat sapien aliquet odio. Integer vitae justo. Aliquam vestibulum fringilla lorem. Sed neque lectus, consectetuer at, consectetuer sed, eleifend ac, lectus. Nulla facilisi (Figure 3). Pellentesque eget lectus. Proin eu metus. Sed porttitor. In hac habitasse platea dictumst. Suspendisse eu lectus. Ut Curabitur tellus magna, porttitor a, commodo a, commodo in, tortor. Donec interdum. Praesent scelerisque. Mae- cenas posuere sodales odio. Vivamus metus lacus, varius quis, imperdiet quis, rhoncus a, turpis. Etiam ligula arcu, elementum a, venenatis quis, sollicitudin sed, metus. Donec nunc pede, tincidunt in, venenatis vitae, faucibus vel.

**TABLE 3** Sideways table caption. For decimal alignment refer column 4 to 9 in tabular* preamble.

| | **Col2 head** | **Col3 head** | **10** | **20** | **30** | **10** | **20** | **30** |
|---|---|---|---|---|---|---|---|---|
| | col2 text | col3 text | 0.7568 | 1.0530 | 1.2642 | 0.9919 | 1.3541 | 1.6108 |
| | | col2 text | 12.5701 | 19.6603 | 25.6809 | 18.0689 | 28.4865 | 37.3011 |
| 3 | col2 text | col3 text | 0.7426 | 1.0393 | 1.2507 | 0.9095 | 1.2524 | 1.4958 |
| | | col3 text | 12.8008 | 19.9620 | 26.0324 | 16.6347 | 26.0843 | 34.0765 |
| | col2 text | col3 text | 0.7285 | 1.0257 | 1.2374 | 0.8195 | 1.1407 | 1.3694* |
| | | col3 text | 13.0360 | 20.2690 | 26.3895 | 15.0812 | 23.4932 | 30.6060† |

*First sideways table footnote. Sideways table footnote. Sideways table footnote. Sideways table footnote.
†Second sideways table footnote. Sideways table footnote. Sideways table footnote. Sideways table footnote.

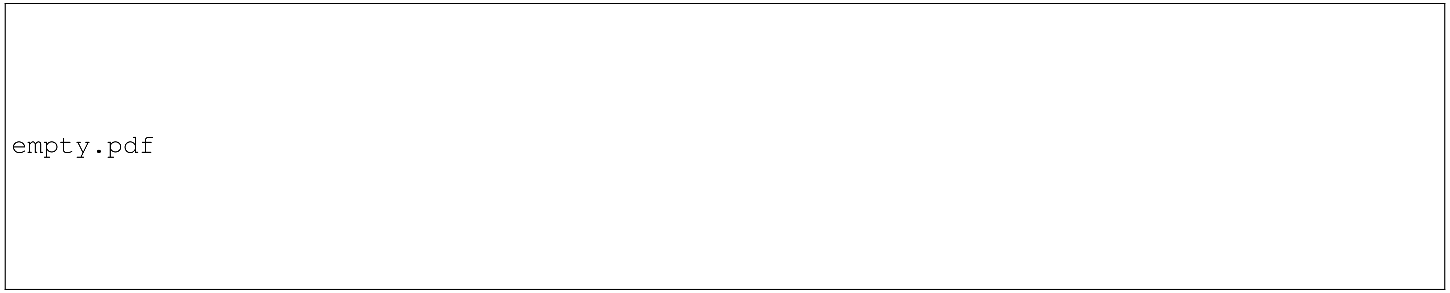


**FIGURE 3** Sideways figure caption. Sideways figure caption. Sideways figure caption. Sideways figure caption. Sideways figure caption. Sideways figure caption.

Pellentesque wisi.[10] Nullam malesuada. Morbi ut tellus ut pede tincidunt porta. Lorem ipsum dolor sit amet, consectetuer adipiscing elit. Etiam congue neque id dolor.

**Algorithm 1** Pseudocode for our algorithm

```
for e doach frame
  for w doater particles $f_i$
    compute fluid flow[1]
    compute fluid-solid interaction[22]
    apply adhesion and surface tension[23]
  end for
  for s doolid particles $s_i$
    for n doeighboring water particles $f_j$
      compute virtual water film
(see Section 3)
    end for
  end for
  for s doolid particles $s_i$
    for n doeighboring water particles $f_j$
      compute growth direction vector
(see Section 2)
    end for
  end for
  for s doolid particles $s_i$
    for n doeighboring water particles $f_j$
      compute $F_\theta$ (see Section 1)
      compute $CE(s_i,f_j)$
(see Section 3)
      if  then$CE(b_i,f_j)$ > glaze threshold
        $j$th water particle's phase ⇐ ICE
      end if
      if  then$CE(c_i,f_j)$ > icicle threshold
        $j$th water particle's phase ⇐ ICE
      end if
    end for
  end for
end for
```

Donec et nisl at wisi luctus bibendum. Nam interdum tellus ac libero. Sed sem justo, laoreet vitae, fringilla at, adipiscing ut, nibh. Maecenas non sem quis tortor eleifend fermentum. Etiam id tortor ac mauris porta vulputate. Integer porta neque vitae massa[1,22]. Maecenas tempus libero a libero posuere dictum. Vestibulum ante ipsum primis in faucibus orci luctus et ultrices posuere cubilia Curae; Aenean quis mauris sed elit commodo placerat. Class aptent taciti sociosqu ad litora torquent per conubia nostra, per inceptos hymenaeos. Vivamus rhoncus tincidunt libero. Etiam elementum pretium justo. Vivamus est. Morbi a tellus eget pede tristique commodo[22]. Nulla nisl. Vestibulum sed nisl eu sapien cursus rutrum.

Pellentesque wisi. Nullam malesuada. Morbi ut tellus ut pede tincidunt porta. Lorem ipsum dolor sit amet, consectetuer adipiscing elit. Etiam congue neque id dolor.

Donec et nisl at wisi luctus bibendum. Nam interdum tellus ac libero. Sed sem justo, laoreet vitae, fringilla at, adipiscing ut, nibh. Integer porta neque vitae massa. Maecenas tempus libero a libero posuere dictum. Vestibulum ante ipsum primis in faucibus orci luctus et ultrices posuere cubilia Curae; Aenean quis mauris sed elit commodo placerat. Maecenas non sem quis tortor eleifend fermentum. Etiam id tortor ac mauris porta vulputate. Integer porta neque vitae massa. Maecenas tempus libero a libero posuere dictum. Vestibulum ante ipsum primis in faucibus orci luctus et ultrices posuere cubilia Curae; Aenean quis mauris sed elit commodo placerat. Class aptent taciti sociosqu ad litora torquent per conubia nostra, per inceptos hymenaeos. Vivamus rhoncus tincidunt libero. Etiam elementum pretium justo. Vivamus est. Morbi a tellus eget pede tristique commodo. Nulla nisl. Vestibulum sed nisl eu sapien cursus rutrum.

$$\begin{aligned}\|\tilde{X}(k)\|^2 &= \frac{\left\|\sum_{i=1}^{p}\tilde{Y}_i(k)+\sum_{j=1}^{q}\tilde{Z}_j(k)\right\|^2}{(p+q)^2}\\ &\le \frac{\sum_{i=1}^{p}\left\|\tilde{Y}_i(k)\right\|^2+\sum_{j=1}^{q}\left\|\tilde{Z}_j(k)\right\|^2}{p+q}.\end{aligned} \tag{1}$$

Sed feugiat. Cum sociis natoque penatibus et magnis dis parturient montes, nascetur ridiculus mus. Ut pellentesque augue sed urna. Vestibulum diam eros, fringilla et, consectetuer eu, nonummy id, sapien. Nullam at lectus. In sagittis ultrices mauris. Curabitur malesuada erat sit amet massa. Fusce blandit. Aliquam erat volutpat. Aliquam euismod. Aenean vel lectus. Nunc imperdiet justo nec dolor.

Etiam euismod. Fusce facilisis lacinia dui. Suspendisse potenti. In mi erat, cursus id, nonummy sed, ullamcorper eget, sapien. Praesent pretium, magna in eleifend egestas, pede pede pretium lorem, quis consectetuer tortor sapien facilisis magna. Mauris quis magna varius nulla scelerisque imperdiet. Aliquam non quam. Aliquam porttitor quam a lacus. Praesent vel arcu ut tortor cursus volutpat. In vitae pede quis diam bibendum placerat. Fusce elementum convallis neque. Sed dolor orci, scelerisque ac, dapibus nec, ultricies ut, mi. Duis nec dui quis leo sagittis commodo.

$$\begin{aligned}\|\tilde{X}(k)\|^2 &= \frac{\left\|\sum_{i=1}^{p}\tilde{Y}_i(k)+\sum_{j=1}^{q}\tilde{Z}_j(k)\right\|^2}{(p+q)^2}\\ &\le \frac{\sum_{i=1}^{p}\left\|\tilde{Y}_i(k)\right\|^2+\sum_{j=1}^{q}\left\|\tilde{Z}_j(k)\right\|^2}{p+q}.\end{aligned} \tag{2}$$

Aliquam lectus. Vivamus leo. Quisque ornare tellus ullamcorper nulla. Mauris porttitor pharetra tortor. Sed fringilla justo sed mauris. Mauris tellus. Sed non leo. Nullam elementum, magna in cursus sodales, augue est scelerisque sapien, venenatis congue nulla arcu et pede. Ut suscipit enim vel sapien. Donec congue. Maecenas urna mi, suscipit in, placerat ut, vestibulum ut, massa. Fusce ultrices nulla et nisl.

Etiam ac leo a risus tristique nonummy. Donec dignissim tincidunt nulla. Vestibulum rhoncus molestie odio. Sed lobortis, justo et pretium lobortis, mauris turpis condimentum augue, nec ultricies nibh arcu pretium enim. Nunc purus neque, placerat id, imperdiet sed, pellentesque nec, nisl. Vestibulum imperdiet neque non sem accumsan laoreet. In hac habitasse platea dictumst. Etiam condimentum facilisis libero. Suspendisse in elit quis nisl aliquam dapibus. Pellentesque auctor sapien. Sed egestas sapien nec lectus. Pellentesque vel dui vel neque bibendum viverra. Aliquam porttitor nisl nec pede. Proin mattis libero vel turpis. Donec rutrum mauris et libero. Proin euismod porta felis. Nam lobortis, metus quis elementum commodo, nunc lectus elementum mauris, eget vulputate ligula tellus eu neque. Vivamus eu dolor.

#### 5 | CONCLUSIONS

Lorem ipsum dolor sit amet, consectetuer adipiscing elit. Ut purus elit, vestibulum ut, placerat ac, adipiscing vitae, felis. Curabitur dictum gravida mauris. Nam arcu libero, nonummy eget, consectetuer id, vulputate a, magna. Donec vehicula augue eu neque. Pellentesque habitant morbi tristique senectus et netus et malesuada fames ac turpis egestas. Mauris ut leo. Cras viverra metus rhoncus sem. Nulla et lectus vestibulum urna fringilla ultrices. Phasellus eu tellus sit amet tortor gravida placerat. Integer sapien est, iaculis in, pretium quis, viverra ac, nunc. Praesent eget sem vel leo ultrices bibendum. Aenean faucibus. Morbi dolor nulla, malesuada eu, pulvinar at, mollis ac, nulla. Curabitur auctor semper nulla. Donec varius orci eget risus. Duis nibh mi, congue eu, accumsan eleifend, sagittis quis, diam. Duis eget orci sit amet orci dignissim rutrum.

Nam dui ligula, fringilla a, euismod sodales, sollicitudin vel, wisi. Morbi auctor lorem non justo. Nam lacus libero, pretium at, lobortis vitae, ultricies et, tellus. Donec aliquet, tortor sed accumsan bibendum, erat ligula aliquet magna, vitae ornare odio metus a mi. Morbi ac orci et nisl hendrerit mollis. Suspendisse ut massa. Cras nec ante. Pellentesque a nulla. Cum sociis natoque penatibus et magnis dis parturient montes, nascetur ridiculus mus. Aliquam tincidunt urna. Nulla ullamcorper vestibulum turpis. Pellentesque cursus luctus mauris.

#### AUTHOR CONTRIBUTIONS

This is an author contribution text. This is an author contribution text. This is an author contribution text. This is an author contribution text. This is an author contribution text.

#### ACKNOWLEDGMENTS

This is acknowledgment text.[24] Provide text here. This is acknowledgment text. Provide text here. This is acknowledgment text. Provide text here. This is acknowledgment text. Provide text here. This is acknowledgment text. Provide text here. This is acknowledgment text. Provide text here. This is acknowledgment text. Provide text here. This is acknowledgment text. Provide text here. This is acknowledgment text. Provide text here.

#### FINANCIAL DISCLOSURE

None reported.

#### CONFLICT OF INTEREST

The authors declare no potential conflict of interests.

#### SUPPORTING INFORMATION

Additional supporting information may be found in the online version of the article at the publisher's website.

**How to cite this article:** Taylor M., Lauritzen P, Erath C, and Mittal R. On simplifying 'incremental remap'-based transport schemes. *J Comput Phys.* 2021;00(00):1–18.

#### APPENDIX

#### A PROGRAM CODES APPEAR IN APPENDIX

Using the package `listings` you can add non-formatted text as you would do with `\begin{verbatim}` but its main aim is to include the source code of any programming language within your document.
Use `\begin{lstlisting}...\end{lstlisting}` for program codes without mathematics.

The `listings` package supports all the most common languages and it is highly customizable. If you just want to write code within your document, the package provides the `lstlisting` environment; the output will be in Computer Modern typewriter font. Refer to the below example:

**LISTING 1** Descriptive caption text

```
for i:=maxint to 0 do
begin
{ do nothing }
end;
Write('Case insensitive ');
WritE('Pascal keywords.');
```

##### A.1 Subsection title of first appendix

Nam dui ligula, fringilla a, euismod sodales, sollicitudin vel, wisi. Morbi auctor lorem non justo. Nam lacus libero, pretium at, lobortis vitae, ultricies et, tellus. Donec aliquet, tortor sed accumsan bibendum, erat ligula aliquet magna, vitae ornare odio metus a mi. Morbi ac orci et nisl hendrerit mollis. Suspendisse ut massa. Cras nec ante. Pellentesque a nulla. Cum sociis natoque penatibus et magnis dis parturient montes, nascetur ridiculus mus. Aliquam tincidunt urna. Nulla ullamcorper vestibulum turpis. Pellentesque cursus luctus mauris.

Nulla malesuada porttitor diam. Donec felis erat, congue non, volutpat at, tincidunt tristique, libero. Vivamus viverra fermentum felis. Donec nonummy pellentesque ante. Phasellus adipiscing semper elit. Proin fermentum massa ac quam. Sed diam turpis, molestie vitae, placerat a, molestie nec, leo. Maecenas lacinia. Nam ipsum ligula, eleifend at, accumsan nec, suscipit a, ipsum. Morbi blandit ligula feugiat magna. Nunc eleifend consequat lorem. Sed lacinia nulla vitae enim. Pellentesque tincidunt purus vel magna. Integer non enim. Praesent euismod nunc eu purus. Donec bibendum quam in tellus. Nullam cursus pulvinar lectus. Donec et mi. Nam vulputate metus eu enim. Vestibulum pellentesque felis eu massa. Nulla malesuada porttitor diam. Donec felis erat, congue non, volutpat at, tincidunt tristique, libero. Vivamus viverra fermentum felis. Donec nonummy pellentesque ante. Phasellus adipiscing semper elit. Proin fermentum massa ac quam. Sed diam turpis, molestie vitae, placerat a, molestie nec, leo. Maecenas lacinia. Nam ipsum ligula, eleifend at, accumsan nec, suscipit a, ipsum. Morbi blandit ligula feugiat magna. Nunc eleifend consequat lorem. Sed lacinia nulla vitae enim. Pellentesque tincidunt purus vel magna. Integer non enim. Praesent euismod nunc eu purus. Donec bibendum quam in tellus. Nullam cursus pulvinar lectus. Donec et mi. Nam vulputate metus eu enim. Vestibulum pellentesque felis eu massa.

##### A.1.1 Subsection title of first appendix

**Unnumbered figure**

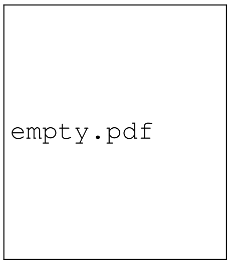


Fusce mauris. Vestibulum luctus nibh at lectus. Sed bibendum, nulla a faucibus semper, leo velit ultricies tellus, ac venenatis arcu wisi vel nisl. Vestibulum diam. Aliquam pellentesque, augue quis sagittis posuere, turpis lacus congue quam, in hendrerit risus eros eget felis. Maecenas eget erat in sapien mattis porttitor. Vestibulum porttitor. Nulla facilisi. Sed a turpis eu lacus commodo facilisis. Morbi fringilla, wisi in dignissim interdum, justo lectus sagittis dui, et vehicula libero dui cursus dui. Mauris tempor ligula sed lacus. Duis cursus enim ut augue. Cras ac magna. Cras nulla.

Nulla egestas. Curabitur a leo. Quisque egestas wisi eget nunc. Nam feugiat lacus vel est. Curabitur consectetuer. Suspendisse vel felis. Ut lorem lorem, interdum eu, tincidunt sit amet, laoreet vitae, arcu. Aenean faucibus pede eu ante. Praesent enim elit, rutrum at, molestie non, nonummy vel, nisl. Ut lectus eros, malesuada sit amet, fermentum eu, sodales cursus, magna. Donec eu purus. Quisque vehicula, urna sed ultricies auctor, pede lorem egestas dui, et convallis elit erat sed nulla. Donec luctus. Curabitur et nunc. Aliquam dolor odio, commodo pretium, ultricies non, pharetra in, velit. Integer arcu est, nonummy in, fermentum faucibus, egestas vel, odio.

#### B SECTION TITLE OF SECOND APPENDIX

Fusce mauris. Vestibulum luctus nibh at lectus. Sed bibendum, nulla a faucibus semper, leo velit ultricies tellus, ac venenatis arcu wisi vel nisl. Vestibulum diam. Aliquam pellentesque, augue quis sagittis posuere, turpis lacus congue quam, in hendrerit risus eros eget felis. Maecenas eget erat in sapien mattis porttitor. Vestibulum porttitor. Nulla facilisi. Sed a turpis eu lacus commodo facilisis. Morbi fringilla, wisi in dignissim interdum, justo lectus sagittis dui, et vehicula libero dui cursus dui. Mauris tempor ligula sed lacus. Duis cursus enim ut augue. Cras ac magna. Cras nulla (Figure B1).

Nulla egestas. Curabitur a leo. Quisque egestas wisi eget nunc. Nam feugiat lacus vel est. Curabitur consectetuer. Suspendisse vel felis. Ut lorem lorem, interdum eu, tincidunt sit amet, laoreet vitae, arcu. Aenean faucibus pede eu ante. Praesent enim elit, rutrum at, molestie non, nonummy vel, nisl. Ut lectus eros, malesuada sit amet, fermentum eu, sodales cursus, magna. Donec eu purus. Quisque vehicula, urna sed ultricies auctor, pede lorem egestas dui, et convallis elit erat sed nulla. Donec luctus. Curabitur et nunc. Aliquam dolor odio, commodo pretium, ultricies non, pharetra in, velit. Integer arcu est, nonummy in, fermentum faucibus, egestas vel, odio.

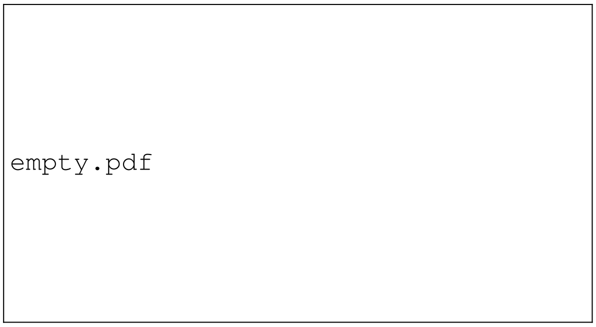


**FIGURE B1** This is an example for appendix figure.

**TABLE B1** This is an example of Appendix table showing food requirements of army, navy and airforce.

| **Col1 head** | **Col2 head** | **Col3 head** | **Col4 head** | **Col5 head** | **Col6 head** |
|---|---|---|---|---|---|
| col1 text | col2 text | col3 text | col4 text | col5 text | col6 text |
| col1 text | col2 text | col3 text | col4 text | col5 text | col6 text |
| col1 text | col2 text | col3 text | col4 text | col5 text | col6 text |

##### B.1 Subsection title of second appendix

Sed commodo posuere pede. Mauris ut est. Ut quis purus. Sed ac odio. Sed vehicula hendrerit sem. Duis non odio. Morbi ut dui. Sed accumsan risus eget odio. In hac habitasse platea dictumst. Pellentesque non elit. Fusce sed justo eu urna porta tincidunt. Mauris felis odio, sollicitudin sed, volutpat a, ornare ac, erat. Morbi quis dolor. Donec pellentesque, erat ac sagittis semper, nunc dui lobortis purus, quis congue purus metus ultricies tellus. Proin et quam. Class aptent taciti sociosqu ad litora torquent per conubia nostra, per inceptos hymenaeos. Praesent sapien turpis, fermentum vel, eleifend faucibus, vehicula eu, lacus.

Pellentesque habitant morbi tristique senectus et netus et malesuada fames ac turpis egestas. Donec odio elit, dictum in, hendrerit sit amet, egestas sed, leo. Praesent feugiat sapien aliquet odio. Integer vitae justo. Aliquam vestibulum fringilla lorem. Sed neque lectus, consectetuer at, consectetuer sed, eleifend ac, lectus. Nulla facilisi. Pellentesque eget lectus. Proin eu metus. Sed porttitor. In hac habitasse platea dictumst. Suspendisse eu lectus. Ut mi mi, lacinia sit amet, placerat et, mollis vitae, dui. Sed ante tellus, tristique ut, iaculis eu, malesuada ac, dui. Mauris nibh leo, facilisis non, adipiscing quis, ultrices a, dui.

###### B.1.1 Subsection title of second appendix

Lorem ipsum dolor sit amet, consectetuer adipiscing elit. Ut purus elit, vestibulum ut, placerat ac, adipiscing vitae, felis. Curabitur dictum gravida mauris. Nam arcu libero, nonummy eget, consectetuer id, vulputate a, magna. Donec vehicula augue eu neque. Pellentesque habitant morbi tristique senectus et netus et malesuada fames ac turpis egestas. Mauris ut leo. Cras viverra metus rhoncus sem. Nulla et lectus vestibulum urna fringilla ultrices. Phasellus eu tellus sit amet tortor gravida placerat. Integer sapien est, iaculis in, pretium quis, viverra ac, nunc. Praesent eget sem vel leo ultrices bibendum. Aenean faucibus. Morbi dolor nulla, malesuada eu, pulvinar at, mollis ac, nulla. Curabitur auctor semper nulla. Donec varius orci eget risus. Duis nibh mi, congue eu, accumsan eleifend, sagittis quis, diam. Duis eget orci sit amet orci dignissim rutrum (Table B1).

Nam dui ligula, fringilla a, euismod sodales, sollicitudin vel, wisi. Morbi auctor lorem non justo. Nam lacus libero, pretium at, lobortis vitae, ultricies et, tellus. Donec aliquet, tortor sed accumsan bibendum, erat ligula aliquet magna, vitae ornare odio metus a mi. Morbi ac orci et nisl hendrerit mollis. Suspendisse ut massa. Cras nec ante. Pellentesque a nulla. Cum sociis natoque penatibus et magnis dis parturient montes, nascetur ridiculus mus. Aliquam tincidunt urna. Nulla ullamcorper vestibulum turpis. Pellentesque cursus luctus mauris.

Example for an equation inside appendix

$$\mathcal{L} = i\bar{\psi}\gamma^{\mu}D_{\mu}\psi - \frac{1}{4}F_{\mu\nu}^{a}F^{a\mu\nu} - m\bar{\psi}\psi \tag{B1}$$

#### C EXAMPLE OF ANOTHER APPENDIX SECTION

This is sample for paragraph text this is sample for paragraph text this is sample for paragraph text this is sample for paragraph text this is sample for paragraph text this is sample for paragraph text this is sample for paragraph text this is sample for paragraph text this is sample for paragraph text this is sample for paragraph text this is sample for paragraph text this is sample for paragraph text this is sample for paragraph text this is sample for paragraph text this is sample for paragraph text this is sample for paragraph text this is sample for paragraph text this is sample for paragraph text this is sample for paragraph text this is sample for paragraph text this is sample for paragraph text this is sample for paragraph text this is sample for paragraph text this is sample for paragraph text this is sample for paragraph text this is sample for paragraph text this is sample for paragraph text this is sample for paragraph text this is sample for paragraph text this is sample for paragraph text this is sample for paragraph text this is sample for paragraph text this is sample for paragraph text this is sample for paragraph text this is sample for paragraph text

Nam dui ligula, fringilla a, euismod sodales, sollicitudin vel, wisi. Morbi auctor lorem non justo. Nam lacus libero, pretium at, lobortis vitae, ultricies et, tellus. Donec aliquet, tortor sed accumsan bibendum, erat ligula aliquet magna, vitae ornare odio metus a mi. Morbi ac orci et nisl hendrerit mollis. Suspendisse ut massa. Cras nec ante. Pellentesque a nulla. Cum sociis natoque penatibus et magnis dis parturient montes, nascetur ridiculus mus. Aliquam tincidunt urna. Nulla ullamcorper vestibulum turpis. Pellentesque cursus luctus mauris.

Nulla malesuada porttitor diam. Donec felis erat, congue non, volutpat at, tincidunt tristique, libero. Vivamus viverra fermentum felis. Donec nonummy pellentesque ante. Phasellus adipiscing semper elit. Proin fermentum massa ac quam. Sed diam turpis, molestie vitae, placerat a, molestie nec, leo. Maecenas lacinia. Nam ipsum ligula, eleifend at, accumsan nec, suscipit a, ipsum. Morbi blandit ligula feugiat magna. Nunc eleifend consequat lorem. Sed lacinia nulla vitae enim. Pellentesque tincidunt purus vel magna. Integer non enim. Praesent euismod nunc eu purus. Donec bibendum quam in tellus. Nullam cursus pulvinar lectus. Donec et mi. Nam vulputate metus eu enim. Vestibulum pellentesque felis eu massa.

$$\mathcal{L} = i\bar{\psi}\gamma^{\mu}D_{\mu}\psi - \frac{1}{4}F^{a}_{\mu\nu}F^{a\mu\nu} - m\bar{\psi}\psi \tag{C2}$$

Nulla malesuada porttitor diam. Donec felis erat, congue non, volutpat at, tincidunt tristique, libero. Vivamus viverra fermentum felis. Donec nonummy pellentesque ante. Phasellus adipiscing semper elit. Proin fermentum massa ac quam. Sed diam turpis, molestie vitae, placerat a, molestie nec, leo. Maecenas lacinia. Nam ipsum ligula, eleifend at, accumsan nec, suscipit a, ipsum. Morbi blandit ligula feugiat magna. Nunc eleifend consequat lorem. Sed lacinia nulla vitae enim. Pellentesque tincidunt purus vel magna. Integer non enim. Praesent euismod nunc eu purus. Donec bibendum quam in tellus. Nullam cursus pulvinar lectus. Donec et mi. Nam vulputate metus eu enim. Vestibulum pellentesque felis eu massa.

Quisque ullamcorper placerat ipsum. Cras nibh. Morbi vel justo vitae lacus tincidunt ultrices. Lorem ipsum dolor sit amet, consectetuer adipiscing elit. In hac habitasse platea dictumst. Integer tempus convallis augue. Etiam facilisis. Nunc elementum fermentum wisi. Aenean placerat. Ut imperdiet, enim sed gravida sollicitudin, felis odio placerat quam, ac pulvinar elit purus eget enim. Nunc vitae tortor. Proin tempus nibh sit amet nisl. Vivamus quis tortor vitae risus porta vehicula.

| **Col1 head** | **Col2 head** | **Col3 head** |
|---|---|---|
| col1 text | col2 text | col3 text |
| col1 text | col2 text | col3 text |
| col1 text | col2 text | col3 text |

Quisque ullamcorper placerat ipsum. Cras nibh. Morbi vel justo vitae lacus tincidunt ultrices. Lorem ipsum dolor sit amet, consectetuer adipiscing elit. In hac habitasse platea dictumst. Integer tempus convallis augue. Etiam facilisis. Nunc elementum fermentum wisi. Aenean placerat. Ut imperdiet, enim sed gravida sollicitudin, felis odio placerat quam, ac pulvinar elit purus eget enim. Nunc vitae tortor. Proin tempus nibh sit amet nisl. Vivamus quis tortor vitae risus porta vehicula.

Fusce mauris. Vestibulum luctus nibh at lectus. Sed bibendum, nulla a faucibus semper, leo velit ultricies tellus, ac venenatis arcu wisi vel nisl. Vestibulum diam. Aliquam pellentesque, augue quis sagittis posuere, turpis lacus congue quam, in hendrerit risus eros eget felis. Maecenas eget erat in sapien mattis porttitor. Vestibulum porttitor. Nulla facilisi. Sed a turpis eu lacus commodo facilisis. Morbi fringilla, wisi in dignissim interdum, justo lectus sagittis dui, evehicula libero dui cursus dui. Mauris tempor ligula sed lacus. Duis cursus enim ut augue. Cras ac magna. Cras nulla. Nulla egestas. Curabitur a leo. Quisque egestas wisi eget nunc. Nam feugiat lacus vel est. Curabitur consectetuer.

Pellentesque habitant morbi tristique senectus et netus et malesuada fames ac turpis egestas. Donec odio elit, dictum in, hendrerit sit amet, egestas sed, leo. Praesent feugiat sapien aliquet odio. Integer vitae justo. Aliquam vestibulum fringilla lorem. Sed neque lectus, consectetuer at, consectetuer sed, eleifend ac, lectus. Nulla facilisi. Pellentesque eget lectus. Proin eu metus. Sed porttitor. In hac habitasse platea dictumst. Suspendisse eu lectus. Ut mi mi, lacinia sit amet, placerat et, mollis vitae, dui. Sed ante tellus, tristique ut, iaculis eu, malesuada ac, dui. Mauris nibh leo, facilisis non, adipiscing quis, ultrices a, dui.

#### AUTHOR BIOGRAPHY

empty.pdf

**Author Name.** Please check with the journal's author guidelines whether author biographies are required. They are usually only included for review-type articles, and typically require photos and brief biographies for each author.
