## Supplementary figures and images for "Inferring Progressive Disconnection in Alzheimer’s Disease with Probabilistic Boolean Networks" (from another version of this preprint: version 2, 2025-07-18)

### binarization_example.png

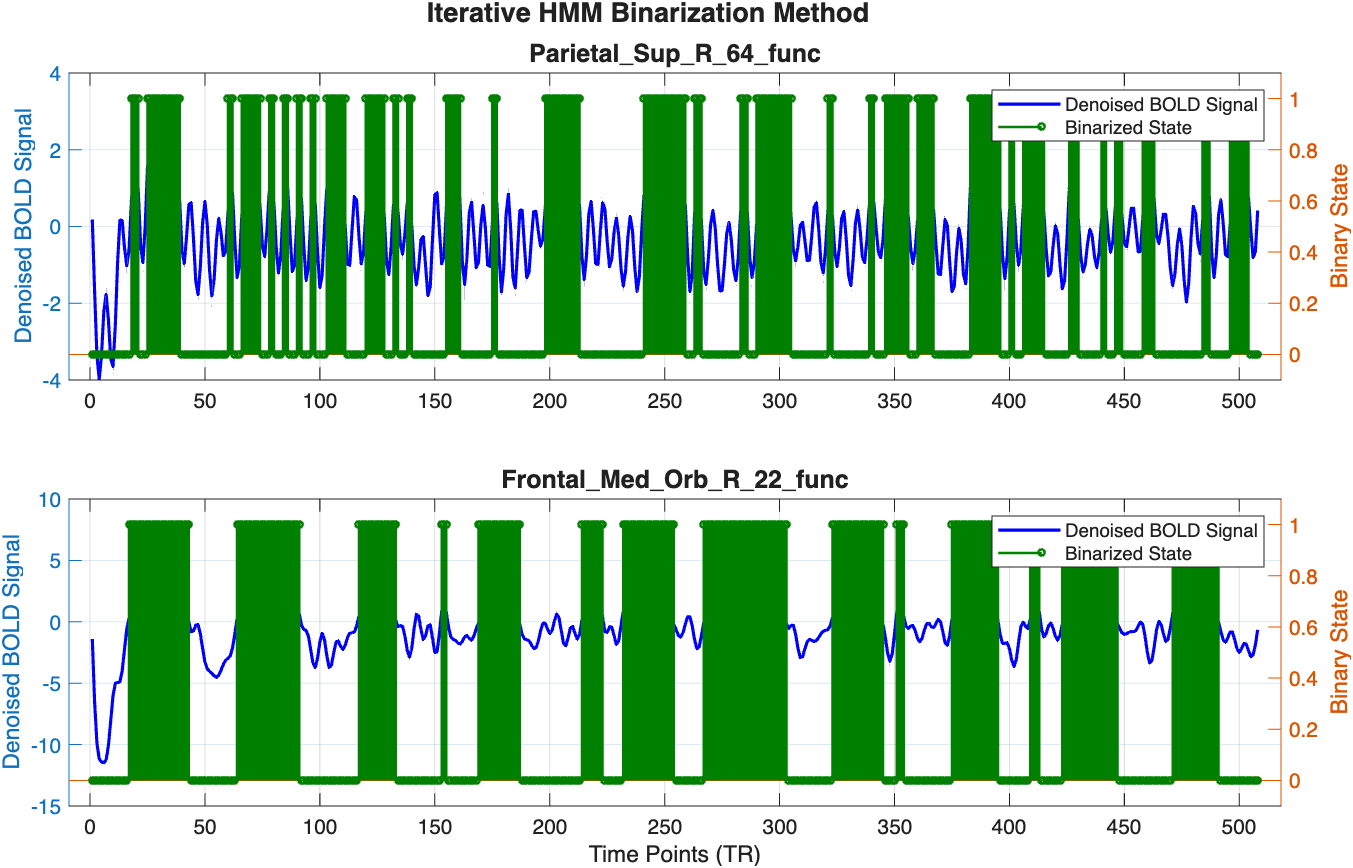

### ci_plot_classic_Frontal_Med_Orb_R_to_Hippocampus_L.png

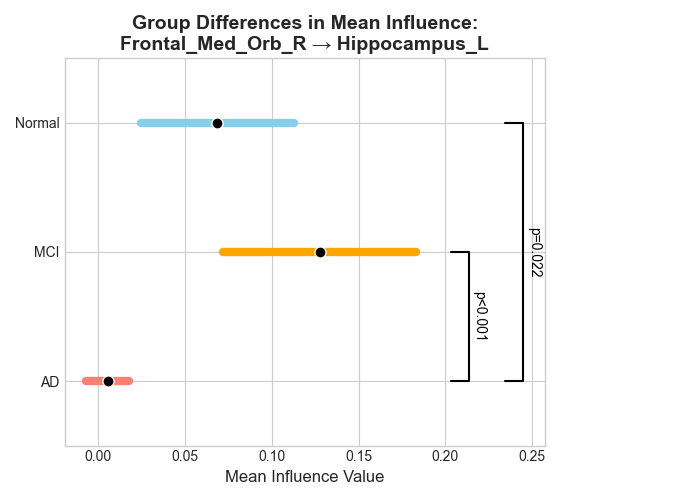

### ci_plot_classic_Precuneus_R_to_Hippocampus_L.png

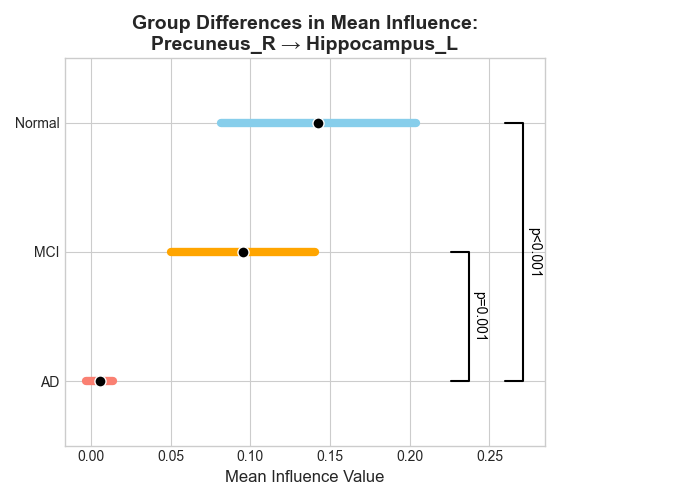

### influence_matrix_graph.png

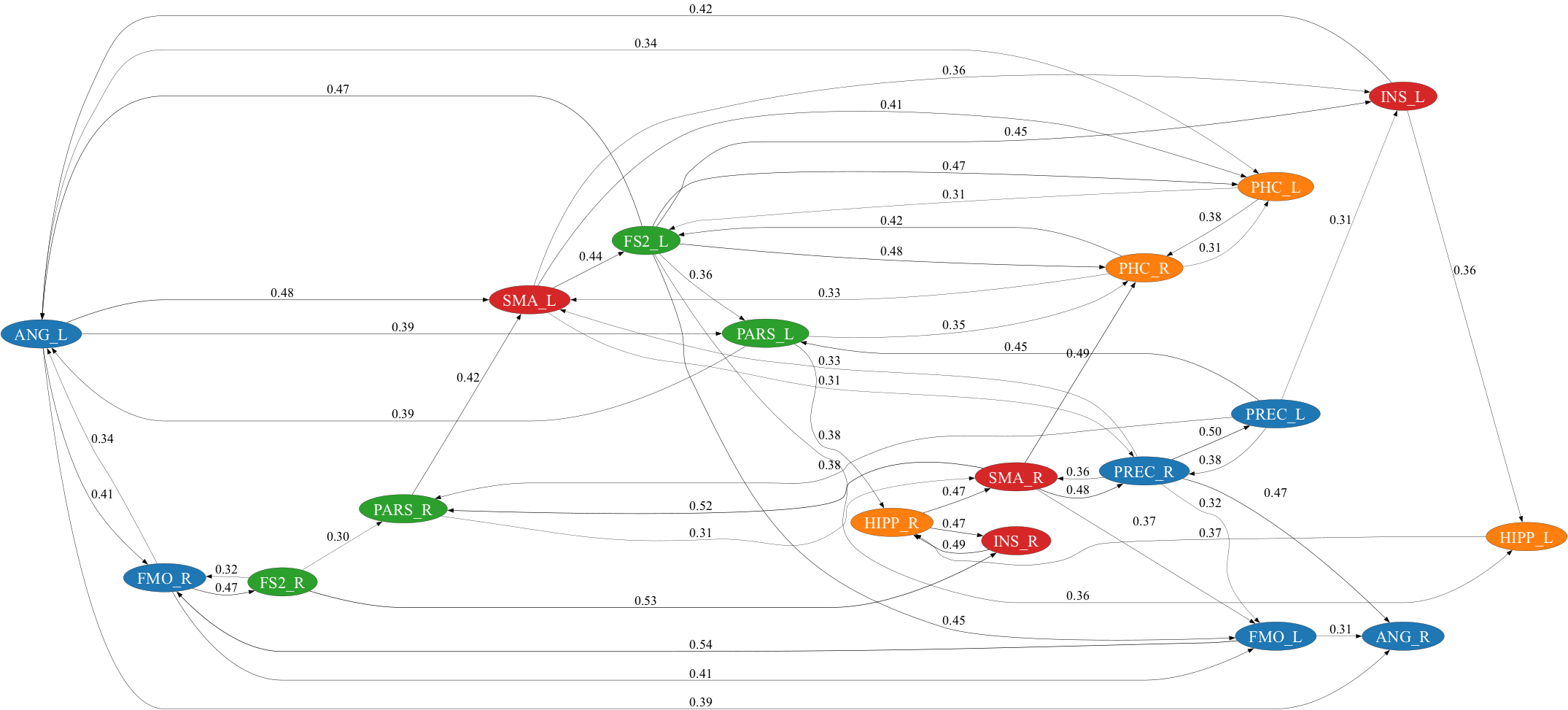

### linear_progression_filtered.png

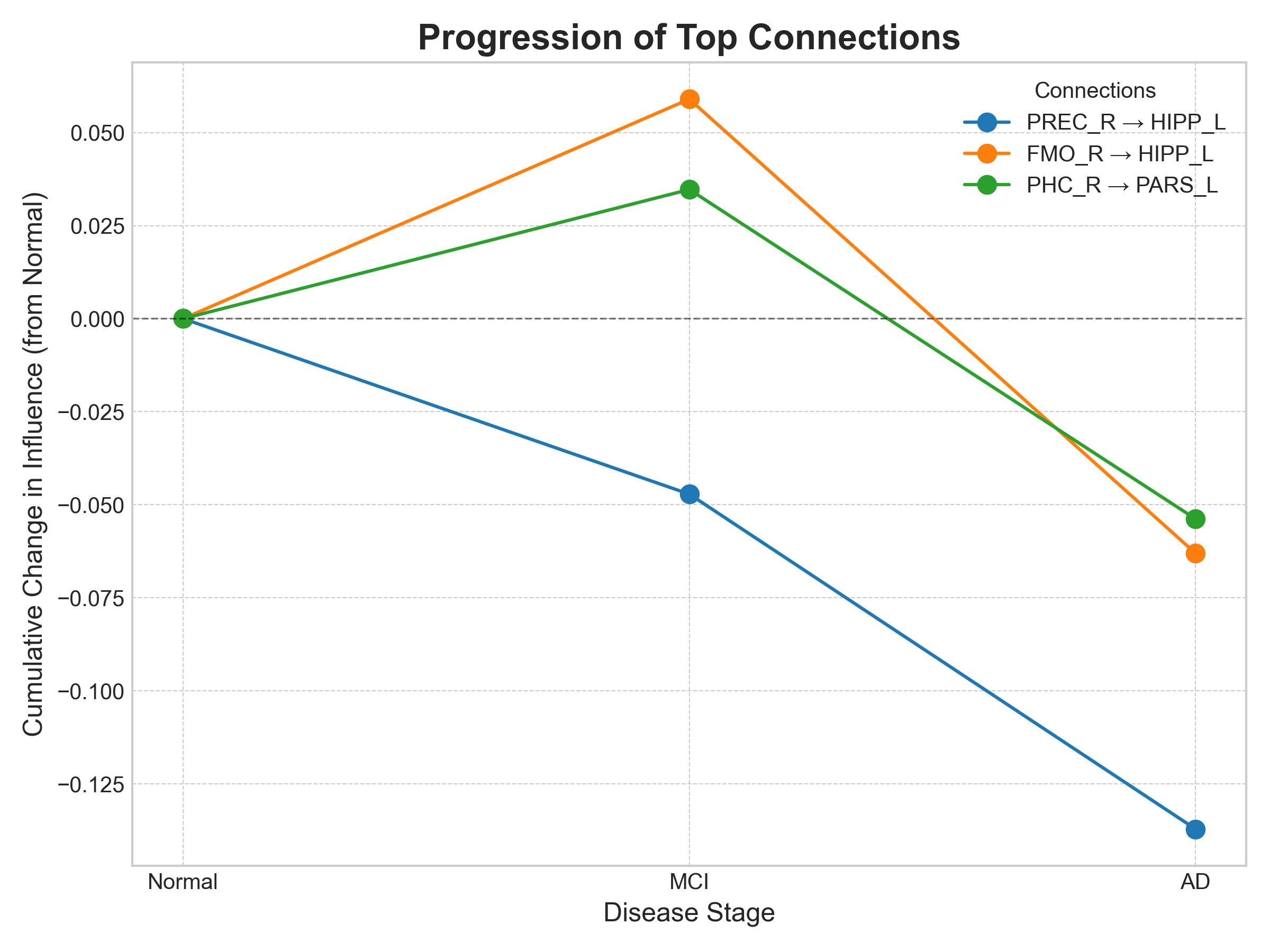

### rhlogo.jpg

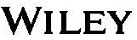

### significant_connections_network.png

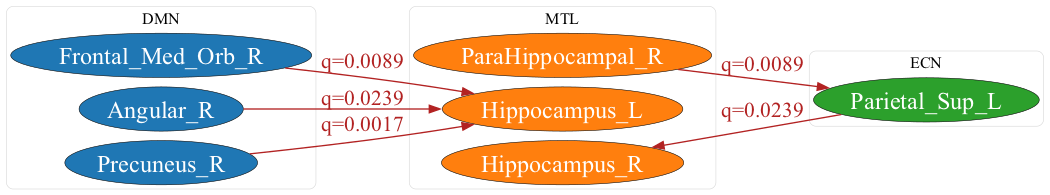
